# *Gβγ* Activates *PIP*2 Hydrolysis by Recruiting and Orienting *PLCβ* on the Membrane Surface

**DOI:** 10.1101/2022.12.20.521270

**Authors:** Maria E. Falzone, Roderick MacKinnon

**Affiliations:** Laboratory of Molecular Neurobiology and Biophysics, The Rockefeller University, New York, United States; Howard Hughes Medical Institute, The Rockefeller University, New York, United States

**Keywords:** *PLCβ*, *Gβγ*, *PIP*2, GPCR signaling, membrane recruitment

## Abstract

*PLCβs* catalyze the hydrolysis of *PIP*2 into IP3 and DAG. *PIP*2 regulates the activity of many membrane proteins, while IP3 and DAG lead to increased intracellular Ca^2+^ levels and activate PKC, respectively. *PLCβs* are regulated by GPCRs through direct interaction with *Gα*_*q*_ and *Gβγ*. This study addresses the mechanism by which *Gβγ* activates *PLCβ*3. We show that *PLCβ*3 functions as a slow Michaelis-Menten enzyme (*k*_*cat*_~2 *sec*^−1^, *K*_*M*_~0.43 *mol*%) on membrane surfaces. Its partition coefficient (*K*_*x*_~2.9 * 10^4^) is such that only a small quantity of *PLCβ*3 exists in the membrane in the absence of *Gβγ*. When *Gβγ* is present, equilibrium binding (*K*_*eq*_~0.009 *mol*%) increases *PLCβ*3 in the membrane, increasing *V*_*max*_ in proportion. Atomic structures on membrane vesicle surfaces show that two *Gβγ* anchor *PLCβ*3 with its catalytic site oriented toward the membrane surface. This principle of activation explains rapid stimulated catalysis with low background catalysis.

## Introduction

Phospholipase C-β (*PLCβ*) enzymes cleave phosphatidylinositol 4,5-bisphosphate (*PIP*2) into inositoltriphosphate (IP3) and diacylglycerol (DAG). Their activity is controlled by G protein coupled receptors (GPCRs) through direct interaction with G proteins (Berridge, 1987; Kadamur and Ross, 2013; Lyon and Tesmer, 2013). Because IP3 increases intracellular calcium, DAG activates protein kinase C (PKC), and levels of *PIP*2 regulate numerous ion channels, the *PLCβ* enzymes under GPCR regulation are central to cellular signaling (Figure 1A) (Hansen, 2015; Poveda et al., 2014; Robinson et al., 2019). There are four *PLCβs* (1-4) in humans; *PLCβ*4 is activated by *Gα*_*q*_ and *PLCβ*1 − 3 are activated by both *Gα*_*q*_ and *Gβγ*. *PLCβ*2/3 are also activated by the small GTPases Rac1/2 (Camps et al., 1992; Hicks et al., 2008; Illenberger et al., 1998; Illenberger et al., 2003; Jezyk et al., 2006; Katz et al., 1992; Smrcka and Sternweis, 1993).

**Figure 1:**
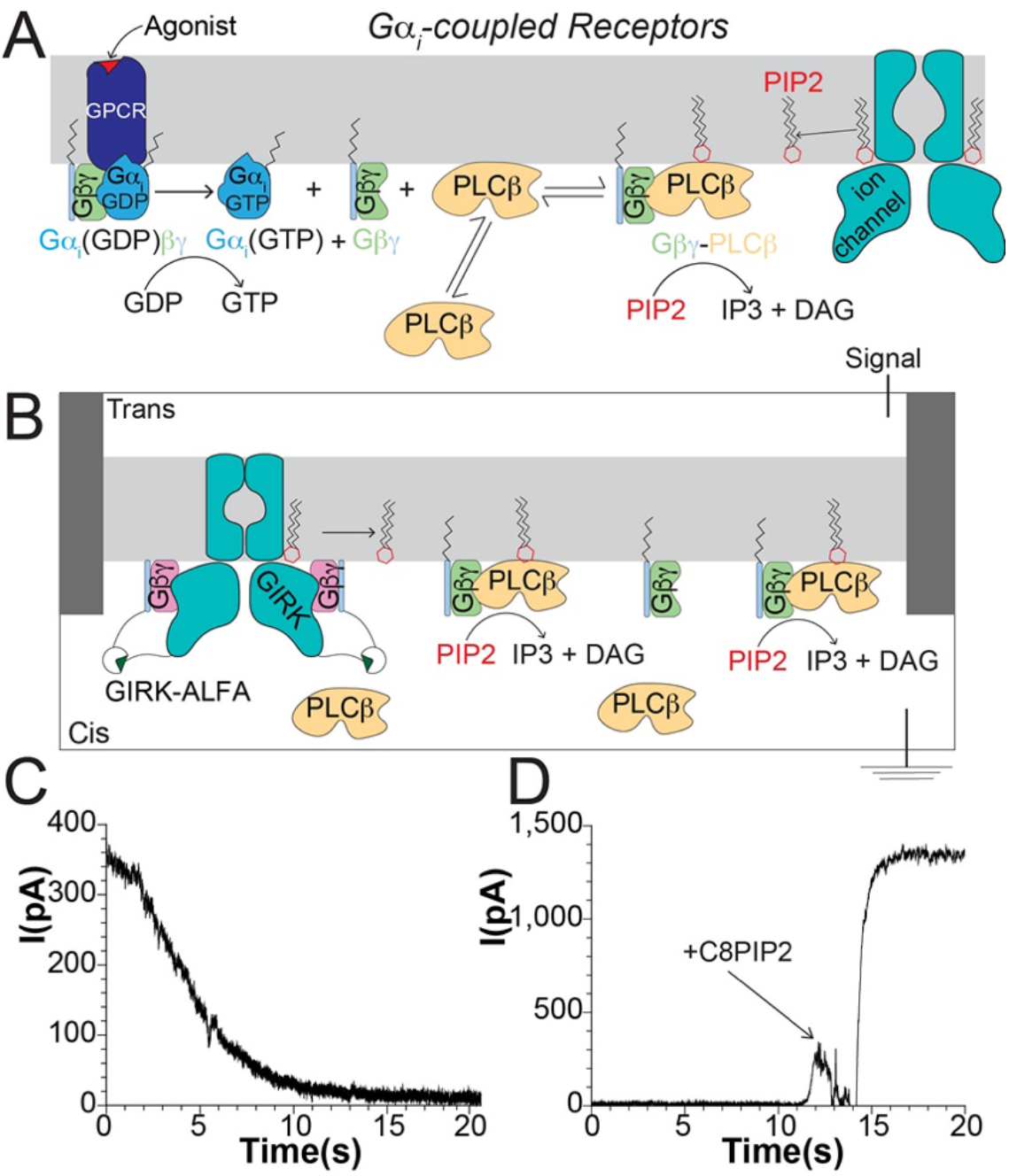
Development of a planar lipid bilayer assay for *PLCβ* activity using a *PIP*2-dependent ion channel as readout. A: Cartoon summary of *Gα*_*i*_-dependent signaling to *PLCβ* through *Gβγ*. B: Cartoon schematic of planar lipid bilayer setup used to measure *PLCβ* function. C: Representative current decay upon *PLCβ*-dependent depletion of *PIP*2. D: Representative current recovery upon reactivation of incorporated channels with short chain C8PIP2. This experiment was carried out under sub-saturating long-chain *PIP*2 (1.0 *mol*%) in the bilayer, which correlates to ~30% of maximal GIRK current. In D, saturating C8PIP2 was added, which leads to ~3x the amount of starting current.

What do we know about *PLCβs* and their regulation by G proteins? *PLCβs* are cytoplasmic enzymes that must access the membrane where their substrate *PIP*2 resides in the inner leaflet. They contain a catalytic core, a proximal C-terminal domain (CTD) with autoinhibitory activity, and a distal CTD with structural homology to a bin-amphiphysin-Rvs (BAR) domain important for membrane binding (Kadamur and Ross, 2013; Lyon and Tesmer, 2013). At the active site, an X-Y linker exerts additional autoinhibitory regulation by direct occlusion (Charpentier et al., 2014; Hicks et al., 2008; Jezyk et al., 2006; Lyon et al., 2014b). *Gα*_*q*_ binds to the proximal and distal CTDs, displacing the autoinhibitory proximal CTD from the catalytic core and Rac1 binds to the PH domain of *PLCβ*2 (Jezyk et al., 2006; Lyon et al., 2014a; Lyon et al., 2013; Lyon et al., 2011; Waldo et al., 2010). Notably, in both cases the autoinhibitory X-Y linker still occludes the active site. Less is known about regulation of *PLCβs* by *Gβγ*. Two potential binding sites have been described, but no structures have been determined (Kadamur and Ross, 2013; Lyon and Tesmer, 2013). The focus of our study is regulation of *PLCβ*3 by *Gβγ*.

The mechanism of *PLCβ* activation by *Gβγ* is unknown. *In vitro* studies have concluded that locally concentrating *PLCβ* on the membrane is not the basis of activation and this has dominated thinking in the field (Jenco et al., 1997; Kadamur and Ross, 2013; Lyon and Tesmer, 2013; Romoser et al., 1996; Runnels et al., 1996). However, the requirement of the lipid group on *Gβγ* to achieve activation and the demonstration that over expression of G proteins in cells increases *PLCβ* in the membrane fraction suggests that a localization mechanism needs revisiting (Gutman et al., 2010; Katz et al., 1992). Part of the challenge in charactering *PLCβ* enzymes is precisely the membrane involvement. *PLCβs* reside in 3 dimensions (the cytoplasm) but catalyze on a 2-dimensional surface (the membrane). Functional measurements must account for this and also permit sufficient time resolution, unlike the standard radioactive assay used in the field until now. To overcome the challenge, we developed new functional methods, including a membrane partitioning assay, rapid kinetic analysis with a direct read-out of *PIP*2 concentration as a function of time, and atomic structures on lipid membrane surfaces, to analyze the mechanism by which *Gβγ* activates *PLCβs*.

## Results

### Development of a planar lipid bilayer assay for *PLCβ3* function

We developed a detergent-free, planar lipid bilayer assay to measure *PLCβ*3 function using a *PIP*2-dependent ion channel to report its concentration over time (Figure 1B-D). Briefly, two chamber cups were connected in the vertical configuration by a ~250 μm hole in a 100 μm piece of Fluorinated ethylene propylene copolymer (Miller, 1983). A ground electrode was placed in the cis chamber and a reference electrode in the trans chamber (Figure 1B). Lipids dispersed in decane were used to paint a bilayer over the hole separating the two chambers. We used a mixture of 2DOPE:1POPC:1POPS lipids and included a pre-determined mole fraction of long-chain *PIP*2 inside the membrane to set its starting concentration. Ion channels and G proteins were incorporated by proteolipid vesicle application to the bilayer and the current from reconstituted ion channels was measured (Miller, 1983). We added *PLCβ*3 to the cis chamber, which was subjected to continuous mixing to ensure homogeneity of the chamber.

The *PIP*2-dependent, G protein-dependent inward rectifying K^+^ channel-2 (GIRK2, specified as GIRK) was used as a readout of *PIP*2 concentration. This channel is well characterized *in vitro*, strictly depends on *PIP*2 for channel opening and is amenable to measuring large macroscopic currents using planar lipid bilayers (Wang et al., 2016; Wang et al., 2014). Further, GIRK exhibits fast rates of association and disassociation of *PIP*2, which permits the measurement of *PLCβ*3 catalytic activity that is not filtered by a slow channel response (Wang et al., 2014). Experiments were carried out in the presence of symmetric MgCl_2_ to ensure blockage of channels with their *PIP*2 binding sites facing the Trans chamber, which is not accessible to *PLCβ*3 (Figure 1B)(Nichols and Lee, 2018). This ensures that when positive voltage is applied, the current is derived only from channels accessible to *PLCβ*3 added to the cis chamber.

GIRK also requires *Gβγ* for channel activity. To separate the effects of *Gβγ* on channel function and *PLCβ*3 activity we used the ALFA nanobody system (Götzke et al., 2019) to tether soluble *Gβγ* to GIRK (Zhao and MacKinnon, 2022). We tagged GIRK with the short ALFA peptide on the C-terminus and *Gγ* with ALFA nanobody on the N-terminus in the background of the C68S mutant, which prevents lipidation of *Gγ*. Nanobody-tagged *Gγ* assembled normally with *Gβ* and was able to bind to other effectors (Zhao and MacKinnon, 2022). Because the ALFA nanobody binds to the ALFA tag with ~30 *pM* affinity (Götzke et al., 2019), at 30 *nM* concentration the ALFA nanobody-tagged *Gβγ* fully activates ALFA peptide-tagged GIRK. In addition, the nanobody-tagged *Gβγ* does not activate *PLCβ*3 due to its lack of a lipid anchor (Katz et al 1992; Gutman et al 2010).

Human *PLCβ*3 was used to establish our assay owing to its significant activation by both *Gβγ* and *Gα*_*q*_ (Philip et al., 2010; Rebres et al., 2011; Smrcka and Sternweis, 1993). The addition of *PLCβ*3 to membranes already containing lipidated *Gβγ*, following an equilibration period of about 2 seconds, led to a rapid current decay that was complete in ~20 s (Figure 1C). Subsequent addition of 32 *μM* C8PIP2 – an aqueous soluble, short chain version of *PIP*2 – rescued the current to a maximum level (Figure 1D)(Wang et al., 2014), indicating that the current decay was due to *PIP*2 depletion from the bilayer by the *PLCβ*3 enzyme. The *PLCβ*3 mediated current decay was slower than when C8PIP2 is rapidly removed by perfusion (Wang et al., 2014). Furthermore, the rate of *PLCβ*3-mediated current decay depends on the *PLCβ*3 concentration (Figure S1). These findings indicate that the decay measures the rate of *PLCβ*3 catalytic activity rather than *PIP*2 unbinding from the channel. No change in the current was observed following *PLCβ*3 addition in the absence of CaCl_2_ (2 mM EGTA), which is required for enzymatic function (Figure S1A). Repetitions of these experiments yielded consistent results with very similar time courses of current decay. These observations indicate that we can measure *PLCβ*3 catalytic activity using this system and that the addition of *PLCβ*3 does not induce artifacts to the bilayer or to reconstituted GIRK channels.

### Kinetic analysis of PIP2 hydrolysis by PLCβ3

The interfacial nature of *PLCβ*3 activity presents a challenge to the study of its function because *PLCβ*3 is a soluble enzyme that must associate with the membrane to carry out catalysis. To describe the reaction occurring at the two-dimensional membrane surface, which must account for the exchange of *PLCβ*3 with the three-dimensional water phase, we give concentrations as dimensionless mole fraction x 100 (*mf*, expressed as *mol*%) using square brackets, [*quantity*], unless specified as molar units using square brackets with subscript *molar*, [*quantity*]_*molar*_. Furthermore, to simplify expressions, we approximate *mf* within each solvent phase, water or lipid, as moles solute per moles solvent rather than moles solute per moles solvent plus solute. This approximation introduces into the kinetic analysis a maximum error in *mf* of 1.0 % for the *PIP*2 concentration in membranes and less than 1.0 % for all other components. For *PIP*2 in membranes, the initial *mf* is pre-determined through the bilayer lipid composition. For *PLCβ*3, the *mf* in membranes is calculated from that in three-dimensional solution using its partition coefficient, which is described below.

The measured current decays can be converted to *PIP*2 decays using the *PIP*2 concentration dependence of the channel, which we determined using titration experiments. Bilayers were formed with varying concentrations of long chain *PIP*2 from 0.1 to 4.0 *mol*%, GIRK-containing vesicles were fused, the current was measured, and water soluble C8PIP2 (32 *μM*) was added to the cis chamber to activate the channels maximally (Figure S1B-C) (Wang et al., 2014). The measured current was normalized to the maximally activated current, *Imax*, for each *PIP*2 concentration and fit to a modified Hill equation, Eq (1), to determine values *A*, *k*, and *r* (Figure 2A):

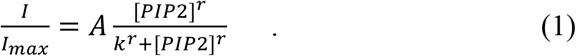

**Figure 2:**
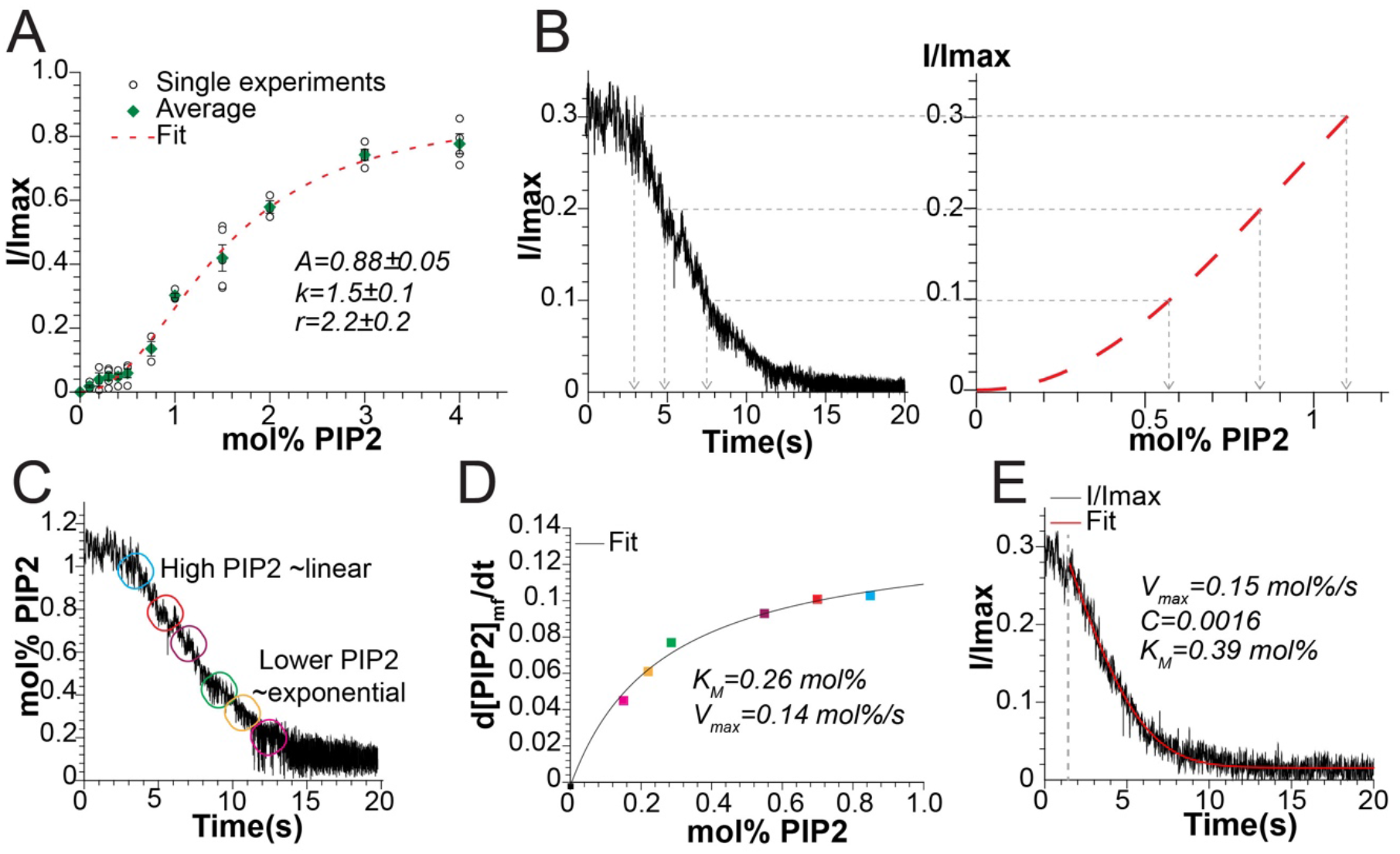
Extraction of values for kinetic parameters for *PLCβ*3 catalysis in the presence of lipidated *Gβγ* from current decay curves A: *PIP*2 activation curve for GIRK varying the *mole*% of *PIP*2 in the bilayer and maximally activating with C8PIP2. Green diamonds are average values, open circles are values from each experiment, and error bars are standard error of the mean. Each point is from 3-5 experiments. The normalized current (I/Imax) is fit to a modified hill equation, Equation 1 (dashed red line). R^2^=0.994. B: Demonstration of using the *PIP*2 activation curve (right) to convert the current decay (left) to *PIP*2 decay. Points on the normalized current decay are matched to *mol*% *PIP*2 and time. C: Resulting *PIP*2 decay over time. Circles denote regions used for integration depicted in D. D: Plot of 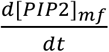 at regions highlights in C vs [*PIP*2] fit to the Michaelis-Menten equation, Equation 2. R^2^=0.993. E: Direct fit (shown as red line) of the normalized current decay to our *PIP*2 activation curve convolved with the time-dependent decay of *PIP*2 with *V*_*max*_ and *K*_*M*_ as free parameters (Equation 4). The gray dashed line denotes where the fit starts which excludes an initial equilibration period. R^2^=0.975.

Eq (1) is an empirical function whose utility is to convert GIRK current into *PIP*2 concentration. In subsequent experiments with *PLCβ*3, bilayers initially contain 1.0 *mol*% *PIP*2, which corresponds to ~30% of the maximal current (Figure 2A).

The *PLCβ*3/*Gβγ*-dependent current decays were corrected by subtracting a constant current value representing non-specific leak, then normalized to the starting *PIP*2 concentration (1.0 *mol*%), and converted to *PIP*2 concentration decays using Eq (1) with the predetermined values for *k*, *A*, and *r* (Figure 2B). After an approximately 2 second delay associated with mixing of *PLCβ*3, *PIP*2 decays contained two components; an initial, approximately linear, component followed by a slower, approximately exponential, component (Figure 2C). The linear component is consistent with *PLCβ*3 catalysis occurring as a 0^th^ order reaction, where the catalytic rate is independent of the *PIP*2 concentration, suggesting that at our starting concentration (1.0 *mol*% *PIP*2) the active site of *PLCβ*3 is nearly fully occupied by substrate (*PIP*2). The second, exponential, component is consistent with the *PIP*2 concentration becoming limiting to catalysis, a first order reaction, as the decay progresses and the concentration of *PIP*2 decreases. In the example shown, for illustrative purpose, we estimated the rate within six intervals along the decay curve, demarcated with different colored circles (Figure 2C), by measuring the slope to approximate 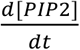 within each interval, and then plotted the slope’s absolute value against the average *PIP*2 concentration for the corresponding interval (Figure 2D). A Michaelis-Menten equation (Eq (2), below) fit the data points with R^2^ ~0.99, indicating that *PLCβ*3 catalytic activity can be described by this kinetic rate equation (Figure 2D).

The graphical procedure described above and in Figure 2 C,D was used as an example to place the *PIP*2 hydrolysis data into a familiar form of rate as a function of substrate concentration. For processing all data, we took a more direct approach to analyze the time dependent decays within the Michaelis-Menten framework. Expressing the Michaelis-Menten rate equation as

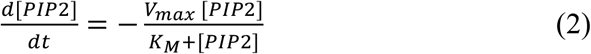

and integrating from t = 0, we obtain for the *PIP*2 concentration as a function of time

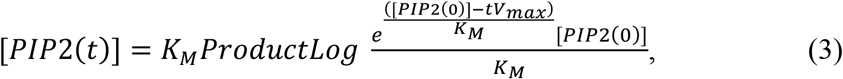

where [*PIP*2(0)] is the *PIP*2 concentration at t = 0 and *K_*w*_* and *V_max_* are the Michaelis-Menten parameters. The *PIP*2 concentration decay is thus given by the Lambert W function, also called the ProductLog function (Goličnik, 2012). Next, substituting Eq (3) into Eq (1), we obtain an expression for GIRK current decay as a function of time due to *PIP*2 hydrolysis,

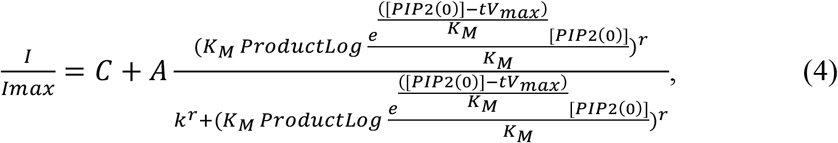

which permits direct fitting of the normalized current decays to estimate *Vmax* and *KM* (Figure 2E). A third free parameter, *C*, accounts for the level of constant background leak (constant current offset) in the bilayer experiments. [*PIP*2(0)], the initial *PIP*2 concentration, is specified by the bilayer composition and *A*, *k* and *r* are predetermined (Figure 2A). This equation fits the current decay data accurately after ~2 seconds (Figure 2E) and yields consistent results for 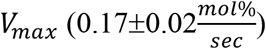 and *K*_*M*_ (0.42±0.05 *mol*%) across repeated experiments (Figure 3C) The value of *C* was typically small and different from one bilayer to the next depending on the level of background leak current.

**Figure 3:**
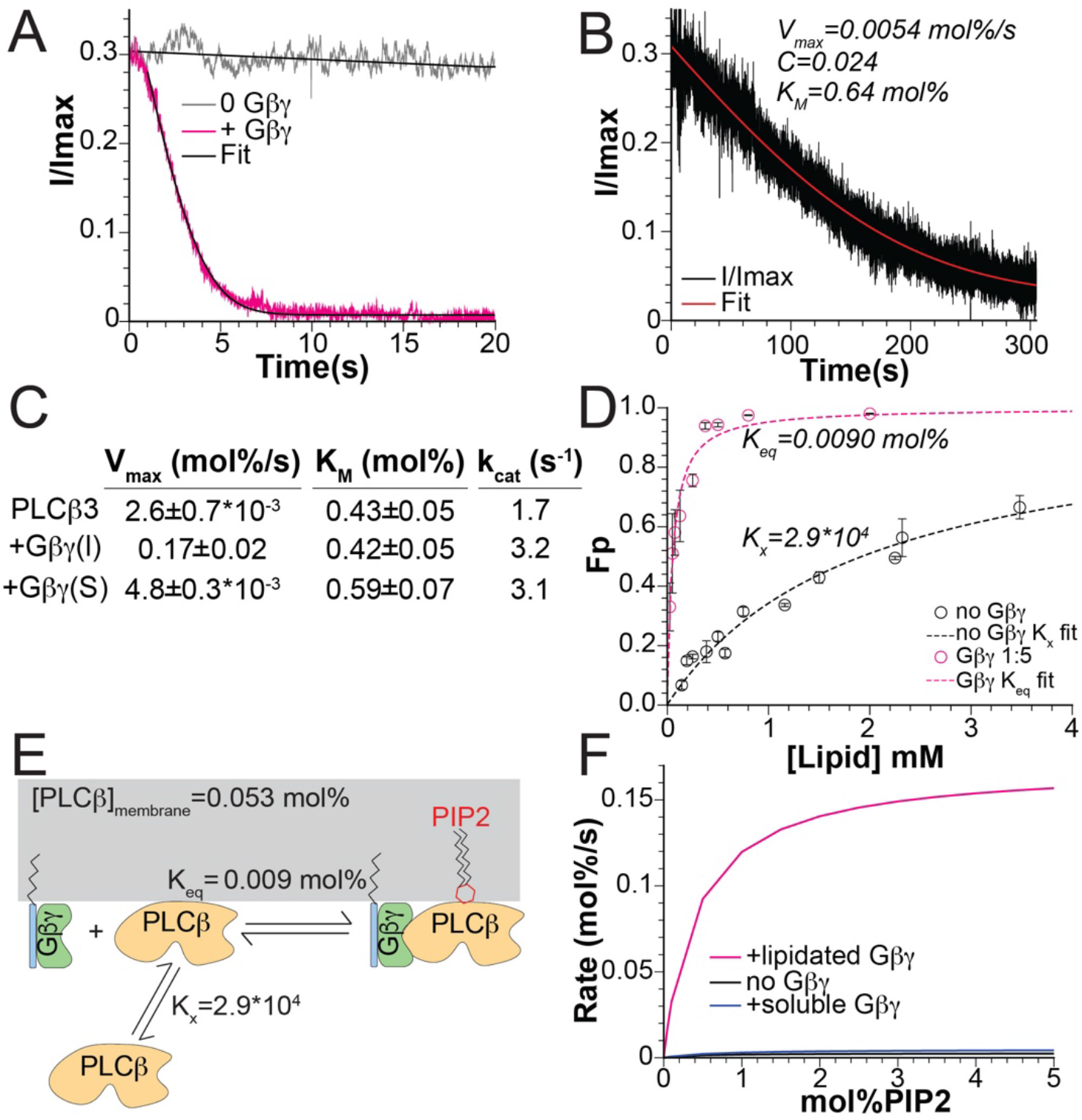
*Gβγ* activates *PLCβ*3 by increasing its concentration at the membrane. A: Comparison of normalized current decay in the presence (pink) and absence (gray) of lipidated *Gβγ* fit to Equation 4 (black lines). Results from the fit without *Gβγ* : 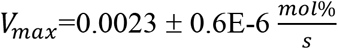, C=−0.03±0.0004, *K*_*M*_=0.43±0.0008 *mol*%, R^2^=0.992. With *Gβγ* : 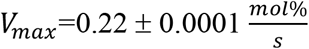, C=0.0074±5E-5, *K*_*M*_=0.37±0.0006 *mol*%. B: Normalized current decay in the presence of 1 *μM* soluble *Gβγ* fit to Equation 4 (red line). R^2^=0.955. C: Comparison of *V*_*max*_, *K*_*M*_, and *k*_*cat*_for *PLCβ*3 alone, with lipidated *Gβγ*, and with soluble *Gβγ*. D: Membrane partitioning curve for *PLCβ*3 alone (black) or in the presence of lipidated *Gβγ* (pink) for 2DOPE:1POPC:1POPS LUVs. Data for 0 *Gβγ* were fit to the Equation 6 for *K*_*x*_ (dashed black lines) and data for +*Gβγ* were fit to Equation 7 for *K*_*eq*_ (White et al., 1998). Error bars are range of mean from two experiments for each lipid concentration. R^2^ =0.962 in the absence of *Gβγ* and R^2^=0.95 in the presence of *Gβγ*. E: Cartoon representation of *PLCβ*3 activation by *Gβγ* through membrane recruitment. *Gβγ* significantly increases the membrane association of *PLCβ*, and accordingly [*PLCβ*]_membrane_, which amplifies *PIP*2 hydrolysis. [*PLCβ*]_membrane_ was calculated using *K*_*x*_ and the solution concentration added in the bilayer experiments (29 nM). *K*_*eq*_ was determined through the fit in panel D. F: Calculated Michaelis-Menten curve for *PLCβ*3 alone (black), in the presence of 1 μM soluble *Gβγ* (blue) or in the presence of lipidated *Gβγ* (pink) using the values for *K*_*M*_ and *V*_*max*_ determined from our fits.

### The role of Gβγ in the function of PLCβ3

In the experiments described above, *Gβγ* was added to the planar lipid bilayers by equilibrating lipid vesicles containing *Gβγ* with the bilayer surface prior to the application of *PLCβ*3. When *Gβγ* is not added to the bilayer, *PLCβ*3 produces a much slower current decay, as shown (Figure 3A, S1D-E). Similarly, in the presence of 1 *μM* aqueous soluble *Gβγ* without a lipid anchor, which does not partition onto the membrane surface (Wang et al., 2014), *PLCβ*3 catalyzed current decay is also slow (Figure 3B,C). Seven experiments were carried out in the absence of *Gβγ* and the RMSD between the current decay curves and Eq (4) were minimized to yield *V*_*max*_ 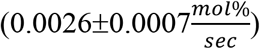 and *K*_*M*_ (0.43±0.05 *mol*%) (Figure 3C). Thus, *Gβγ* in the membrane increases *V*_*max*_ ~65-fold without affecting *K*_*M*_ (Figure 3C).

Because *PLCβ*3 is soluble in aqueous solution but must localize to the membrane surface to catalyze *PIP*2 hydrolysis, we next examined whether *Gβγ* in the membrane influences *PLCβ*3 membrane localization. As detailed by White and colleagues, protein association with membranes cannot be considered as a simple binding equilibrium due to the fluid nature of the membrane without discrete binding sites (White et al., 1998). Instead, membrane association must be treated as a partitioning process between two immiscible solvents, the membrane and the aqueous solution. The equilibrium partition coefficient, *K_x_,* is the ratio of the mole fraction of *PLCβ*3 in the membrane (subscript *m*) to that in aqueous solution (subscript *w*) (White et al., 1998),

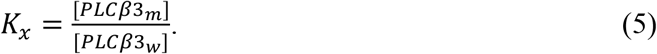

To determine the value of *K*_*x*_, detergent-free liposomes were reconstituted using 2DOPE:1POPC:1POPS lipids to match the lipid composition of the bilayer experiments, and H^+^ nuclear magnetic resonance (NMR) was used to measure the lipid concentration at the end of the detergent removal process (Figure S2A). Large unilamellar vesicles (LUVs) were prepared from the reconstituted liposomes using freeze-thaw cycles and extrusion through a 200 *nm* membrane. The LUVs were incubated with *PLCβ*3 and pelleted using ultracentrifugation to separate the membrane-bound and aqueous protein fractions. This method allows direct measurement of both the bound and free protein using fluorescently labeled *PLCβ*3, which facilitates determining the partition coefficient from each experiment individually (White et al., 1998). The membrane associated fraction of *PLCβ*3 is

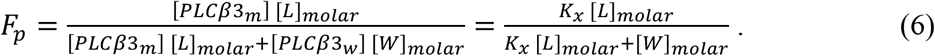

[*W*]_*molar*_, the molar concentration of water is ~ 55 *M* and [*L*]_*molar*_, the molar concentration of lipid, is set for each experiment using a stock measured by NMR. Thus, Eq (6) is a function of the single free parameter, *K*_*x*_, which we determine by fitting Eq (6) to the partitioning data, yielding *K*_*x*_ ~2.9 ∙ 10^4^ (Figure 3D, black curve).

LUVs with the same lipid composition were also prepared containing *Gβγ*, which is exclusively membrane bound, at a protein to lipid ratio of 1:5 (wt:wt), corresponding to 0.34 *mol*%, to match the concentration of *Gβγ* in vesicles equilibrated with planar lipid bilayers in the kinetic experiments. At this concentration of *Gβγ*, we observe that *PLCβ*3 binds to vesicles much more readily than in the absence of *Gβγ* (Figure 3D). This observation is explicable if, when *PLCβ*3 partitions onto the membrane surface, it binds to *Gβγ*. Writing the binding reaction on the membrane surface as *PLCβ*3_*m*_ + *Gβγ* ⇌ *PLCβ*3 ∙ *Gβγ*, we have 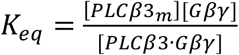 (Figure 3E). (Note that subscript *m* indicates *PLCβ*3 on the membrane. Since *Gβγ* only resides on the membrane a subscript is not used for [*Gβγ*] and [*PLCβ*3 ∙ *Gβγ*].) When equilibrium is reached, the membrane surface will contain a quantity of *PLCβ*3 in the membrane that is not bound to *Gβγ*, set by *K*_*x*_ and the aqueous solution concentration of *PLCβ*3, as well as a quantity of *PLCβ*3 in the membrane that is bound to *Gβγ* (i.e., *PLCβ*3 ∙ *Gβγ*), set by the membrane concentrations of *PLCβ*3, *Gβγ* and *K*_*eq*_. Therefore, in the presence of a total quantity of *Gβγ* on the membrane, [*Gtot*] = [*Gβγ*] + [*Gβγ* ∙ *PLCβ*], the fraction of *PLCβ*3 on the membrane surface, unbound plus bound to *Gβγ*, is given by (see Supplemental Item 1)

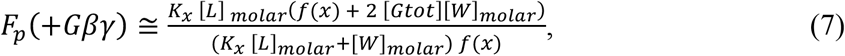

with *f*(*x*) = *p* + *q* + *x* + {4 *q x* + (*p* − *q* + *x*)^2^}^1/2^, where *p* = *K*_*x*_ [*L*]_*molar*_ [*G*_*tot*_], *q* = *K*_*x*_ [*PLC*_*tot*_]_*molar*_, and *x* = *K*_*eq*_ ([*W*]_*molar*_ + *K*_*x*_ [*L*]_*molar*_). Because [*PLC*_*tot*_]_*molar*_ (the molar concentration of *PLCβ*3 (*PLCβ*3_*w*_ and *PLCβ*3_*m*_) plus *PLCβ*3 ∙ *Gβγ*), [*L*]_*molar*_ and [*W*]_*molar*_ (molar concentrations of lipid and water) and [*Gtot*] (*mf Gβγ* plus *PLCβ*3 ∙ *Gβγ* in the membrane) are established in the experimental set up, and *K*_*x*_ is determined through partition measurements in the absence of *Gβγ* (Figure 3D), the right-hand side of Eq (7) contains a single free parameter, *K*_*eq*_, for the binding of *PLCβ*3 to *Gβγ* on the lipid membrane surface. The red dashed curve in Figure 3D corresponds to *K*_*eq*_ = 0.0090 *mol*%. It may seem at first surprising that the series of partitioning experiments in the presence of *Gβγ*, with knowledge of *K*_*x*_ for *PLCβ*3 in the absence of *Gβγ*, uncovers the equilibrium reaction between *PLCβ*3 and *Gβγ* on the membrane surface. But the binding reaction is discernable by this approach, and the inescapable conclusion is that *Gβγ* concentrates *PLCβ*3 on the membrane surface (Figure 3E).

The *PLCβ*3-concentrating effect of *Gβγ* has obvious implications for interpreting the kinetic data reported above, which show that *Gβγ* increases *V*_*max*_ by a factor ~65, without affecting *K*_*M*_ very much (Figure 3C). From Eq (2), *V*_*max*_ is the asymptotic rate of *PIP*2 hydrolysis when [*PIP*2] far exceeds *K*_*M*_. In this limit, the maximum rate of hydrolysis, *V*_*max*_, is given by the total membrane concentration of *PLCβ*3 times *k*_*cat*_, the turnover rate of a *PLCβ*3 ∙ *PIP*2 complex. In the bilayer chamber used for the kinetic experiments, the volume of the aqueous solution is so large compared to the small area of the lipid bilayer that surface binding does not significantly alter [*PLCβ*3_*w*_]. Under this condition, we have

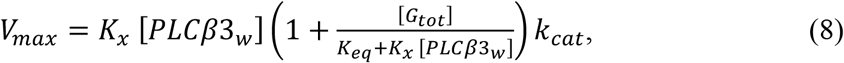

where *K*_*x*_ [*PLCβ*3_*w*_] is the membrane concentration of *PLCβ*3 in the absence of *Gβγ* ([*G*_*tot*_] = 0) and 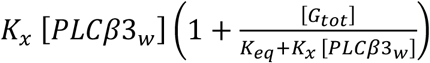 is the membrane concentration in its presence ([*G*_*tot*_] > 0). Thus, 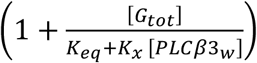 is a multiplier giving the fold-increase in total membrane *PLCβ*3 concentration due to the presence of *Gβγ* at concentration [*G*_*tot*_]. When the known quantities are entered for our experimental conditions, this factor is ~33. In the kinetic experiments we observed a 65-fold increase in *V*_*max*_ in the presence of *Gβγ*. Eq (8) predicts a 33-fold increase through *Gβγ*′*s* ability to increase the local concentration of *PLCβ*3 on the membrane surface. A mere 2-fold increase in *k*_*cat*_ produced by *Gβγ* binding to *PLCβ*3 would account for the full enhancement of *V*_*max*_ in the kinetic experiments (Figure 3C). The important conclusion is that most of the increase in *V*_*max*_ (within a factor of ~2) is explained by the ability of *Gβγ* to concentrate *PLCβ*3 on the membrane surface. Using a conventional Michaelis-Menten plot, with the *V*_*max*_ and *K*_*M*_ values derived experimentally, we observe that at concentrations in our assay, *Gβγ* essentially switches the *PLCβ*3 enzyme on (Figure 3F), and this effect is due largely to the ability of *Gβγ* to concentrate *PLCβ*3 on the membrane surface.

In summary, the kinetic studies show that *PLCβ*3 catalyzes PIP2 hydrolysis with a substrate concentration dependence like that of a Michaelis-Menten enzyme (Figure 2C-E). We note that *K*_*M*_ corresponds to the mid-range of known *PIP*2 concentrations in cell membranes (Figure 2D, 3F) (Mandal, 2020; McLaughlin et al., 2002). *PLCβ*3 aqueous-membrane partition studies show that *Gβγ* concentrates *PLCβ*3 on the membrane surface, enough to account for most of the effect on *V*_*max*_ (Figure 3C, 3D). To a smaller extent (~ 2-fold), *Gβγ* augments *V*_*max*_ through *k*_*cat*_ (Figure 3C). Next, we evaluate the structural underpinnings of these functional properties.

### Structural studies of PLCβ3 in aqueous solution by cryo-EM

We next determined the structure of *PLCβ*3 in aqueous solution using cryo-EM. The structure, consisting of the *PLCβ*3 catalytic core at 3.6 Å resolution, contained the PH domain, EF hands, X and Y domains, the C-terminal part of the X-Y linker, the C2 domain, and the active site with a Ca^2+^ ion bound (Figure 4A-B, S3, Table S1). The autoinhibitory Hα2’ element in the proximal CTD was also resolved, bound to the catalytic core between the Y domain and the C2 domain, as proposed by Lyon and colleagues (Figure 4A-B) (Lyon et al., 2014b; Lyon et al., 2011), but not the distal CTD. We also obtained several low-resolution reconstructions with varying levels of density corresponding to the catalytic core and distal CTD with differing arrangements between the two domains (Figure S3). This observation suggests that the distal CTD is disordered rather than proteolyzed in our final reconstruction and that the two domains are flexible with respect to each other, as previously proposed (Lyon et al., 2013). The catalytic core resolved by cryo-EM is very similar to the crystal structure with a Cα RMSD of 0.6 Å if the Hα2’ helix is excluded (Figure 4B). We note that, as in the crystal structure, the autoinhibitory X-Y linker occludes the active site (Figure 4B). We attempted to determine a structure of *PLCβ*3 in complex with *Gβγ* in solution, in the presence or absence of detergent, without success. Furthermore, we were unable to detect the formation of a complex in solution by size-exclusion chromatography (Figure S3E).

**Figure 4:**
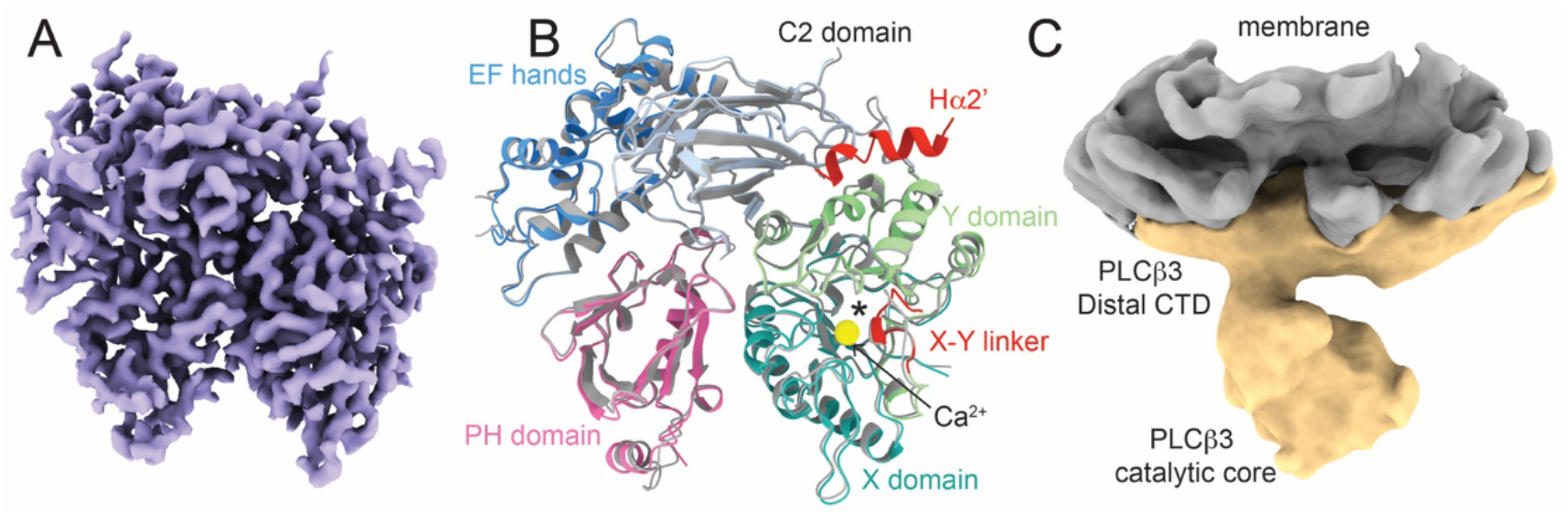
Structures of *PLCβ*3 in solution and on vesicles with and without *Gβγ*. A: Sharpened, masked map of *PLCβ*3 catalytic core obtained from a sample in solution without membranes or detergent. B: Structural alignment of the catalytic core of *PLCβ*3 from the crystal structure of the full-length protein bound to *Gα*_*q*_ (colored in gray, PDBID: 4GNK, (Lyon et al., 2013)) and the structure determined using cryo-EM without membranes (colored by domain). Cα RMSD is 0.6 Å. Calcium ion from the cryo-EM structure is shown as a yellow sphere and the active site is denoted with an asterisk. The PH domain is pink, the EF hand repeats are blue, the C2 domain is light blue, the Y domain is green, the X domain is teal, the X-Y linker and the Hα2’ are red. C: Unsharpened reconstruction of *PLCβ*3 bound to lipid vesicles containing 2DOPE:1POPC:1POPS. *PLCβ*3 is colored in yellow and the membrane is colored in gray.

### Structural studies of PLCβ3 associated with liposomes

We next determined the structure of *PLCβ*3 bound to liposomes consisting of 2DOPE:1POPC:1POPS. We obtained a low-resolution reconstruction with the distal CTD associated with the membrane and the catalytic core located away from the membrane surface (Figure 4C, S4, Table S1). Although the map was low resolution, previously determined structures fit into the density for each domain and all reconstructions showed the same orientation of the protein on the membrane surface (Figure S4). The position of the catalytic core indicates that significant rearrangements of *PLCβ*3 with respect to the membrane must be involved in activation, because the active site is too far from the membrane to access *PIP*2. Activating rearrangements could be mediated by interactions of lipid anchored G proteins with the *PLCβ*3 catalytic core.

### The *PLCβ3/Gβγ* complex on liposomes reveals two *Gβγ* binding sites

We reconstituted *Gβγ* into liposomes consisting of 2DOPE:1POPC:1POPS at a protein to lipid ratio of 1:15 (wt:wt) and incubated the liposomes with purified *PLCβ*3 prior to grid preparation. We determined the structure of the *PLCβ*3 ∙ *Gβγ* complex to 3.5 Å and observed two *Gβγs* bound to the catalytic core of *PLCβ*3 (Figure 5A-C, S5, Table S1). The distal CTD is not resolved in our reconstructions, suggesting that it might adopt many different orientations on the plane of the membrane relative to the catalytic core, in agreement with previous studies showing that heterogeneity in the distal CTD increases upon *Gβγ* binding (Fisher et al., 2020). The catalytic core is very similar to our cryo-EM structure without membranes, with a Cα RMSD of 0.7 Å. Only small rearrangements occur at the *Gβγ* binding sites (Figure S6A). Both autoinhibitory elements, the Hα2’ and the X-Y linker, are engaged with the catalytic core (Figure 5C, S6A) consistent with previous proposals that *Gβγ* does not play a role in relieving this autoinhibition (Hicks et al., 2008; Lyon et al., 2014b; Lyon et al., 2011).

**Figure 5:**
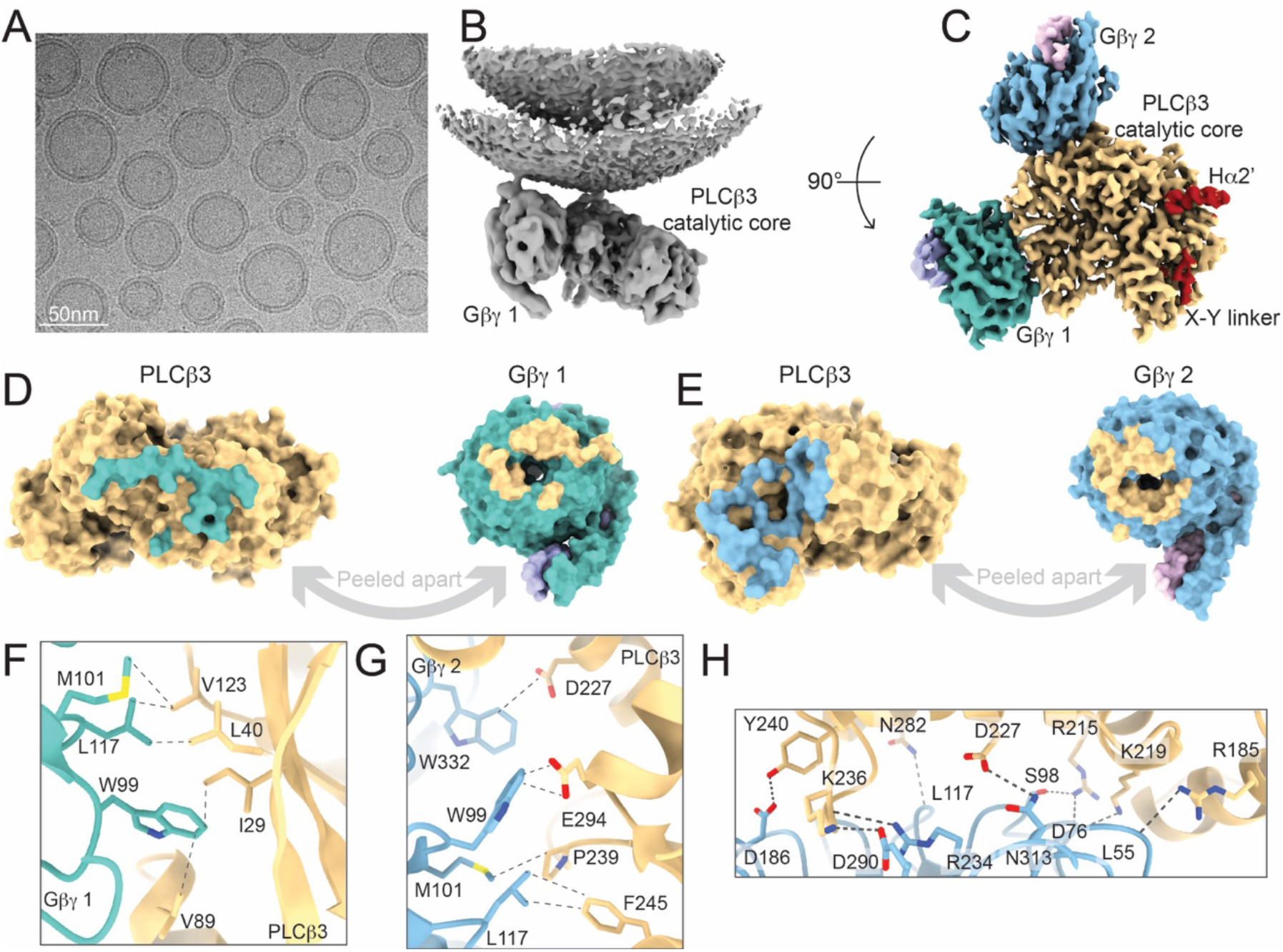
*PLCβ*3 ∙ *Gβγ* complex on lipid vesicles and *Gβγ* interfaces. A: Example micrograph showing lipid vesicles with protein complexes. B: Unsharpened map from non-uniform refinement showing the *PLCβ*3 ∙ *Gβγ* complex on the vesicle surface. C: Sharpened, masked map of the catalytic core of *PLCβ*3 in complex with two *Gβγ*s on lipid vesicles containing 2DOPE:1POPC:1POPS. *PLCβ*3 is yellow, *Gβ* 1 is dark teal, *Gγ* 1 is light purple, *Gβ* 2 is light blue and *Gγ* 2 is light pink. The autoinhibitory elements Hα2’ and the X-Y linker are colored in red. Coloring is the same throughout. D-E: Surface representation of the *PLCβ*-*Gβγ* 1 (D) or *PLCβ*-*Gβγ* 2 (E) interfaces peeled apart to show extensive interactions. Residues on *PLCβ*3 that interact with *Gβγ* 1 or 2 are colored according to the corresponding *Gβ* coloring and residues on the *Gβ*s that interact with *PLCβ*3 are colored in gray. Interface residues were determined using the ChimeraX interface feature using a buried surface area cutoff of 15 Å^2^. F-G: Interactions of residues on *Gβ* that have been shown to be important for *PLCβ* activation with residues from *PLCβ*3 in the *PLCβ*3-*Gβγ* 1 interface (F) or the *PLCβ*3-*Gβγ* 2 interface (G) (Ford et al., 1998). All labeled interactions are < ~4 Å. Interacting residues are shown as sticks and colored by heteroatom. Interactions are denoted by black dashed line. H: Extensive hydrogen bond network in the *PLCβ*3-*Gβγ* 2 interface including both sidechain and backbone interactions. All labeled hydrogen bonds are between ~2.3 and ~3.7 Å. Interacting residues are shown as sticks and colored by heteroatom. Interactions are denoted by black dashed line.

One *Gβγ* is bound to the PH domain and the first EF hand, referred to as *Gβγ* 1, and the other is bound to the remaining EF hands, referred to as *Gβγ* 2 (Figure 5C, S6A-B). Both interfaces are extensive, with the *Gβγ* 1 interface burying ~800 Å^2^ and involving 34 residues, (16 from *PLCβ*3 and 18 from *Gβγ*) and the *Gβγ* 2 interface burying ~1100 Å^2^ and involving 44 residues (21 on *PLCβ*3 and 23 on *Gβγ*) (Figure 5D-E, S6B). The *Gβγ* 1 interface is mostly comprised of hydrophobic interactions, with three hydrogen bonds (Figure 5F, S6C), whereas the *Gβγ* 2 interface is mostly comprised of electrostatic interactions, including nine hydrogen bonds spanning the length of the interface (Figure 5G-H). Both interfaces involve the same region of *Gβγ* that interacts with *Gα* and several residues on Gβ shown to be important for *PLCβ* activation are involved (Ford et al., 1998) (Figure 5E-F). Specifically, M101, L117, and W99 on Gβ 1 form hydrophobic interactions with V123, L40, I29, and V89 on PLCβ3 (Figure 5F, Table S2) (Ford et al., 1998). On Gβ 2, W332 and W99 form electrostatic interactions with D227 and E294 on *PLCβ*3, M101 and L117 form hydrophobic interactions with P239 and F245 on *PLCβ*3, and D186 forms a hydrogen bond with Y240 on *PLCβ*3 (Figure 5G-H, Table S2) (Ford et al., 1998).

We also determined the structure of the *PLCβ*3 ∙ *Gβγ* complex using lipid nanodiscs. We reconstituted *Gβγ* into nanodiscs formed using the MSP2N2 scaffold protein(Grinkova et al., 2010) and 2DOPE:1POPC:1POPS lipids and incubated them with purified *PLCβ*3 prior to grid preparation. We observed only reconstructions with 2 *Gβγs* bound and determined the structure of the complex to 3.3 Å (Figure S7). The two *Gβγ* are bound in the same locations as was observed in liposomes with comparable interfaces (Figure S7C). A model for this structure aligns well to the model built using the lipid vesicle reconstruction with a Cα RMSD of 0.8 Å for all proteins (Figure S7E). These structures suggest that the *PLCβ*3 ∙ *Gβγ* complex depends on a membrane environment as we were unable to form a stable complex in solution with or without detergent, which highlights the importance of the membrane in *Gβγ* −dependent activation of *PLCβ*3.

### *Gβγ* mediates membrane association and orientation of the *PLCβ3* catalytic core

Unmasked refinement of our final subset of particles from the liposome structure yielded a 3.8 Å reconstruction showing the *PLCβ*3 ∙ *Gβγ* assembly and density from the membrane (Figure 5B, 6A). The two *Gβγs* and the region of *PLCβ*3 between them are closely associated with the membrane and the remainder of the catalytic core, including the active site, tilts away from the membrane (Figure 6A). Despite the tilting, the structure reveals significant rearrangement of the catalytic core with respect to the membrane compared to its position in the absence of *Gβγ*, where it was separated from the membrane surface by a larger distance (Figure 6A).

**Figure 6:**
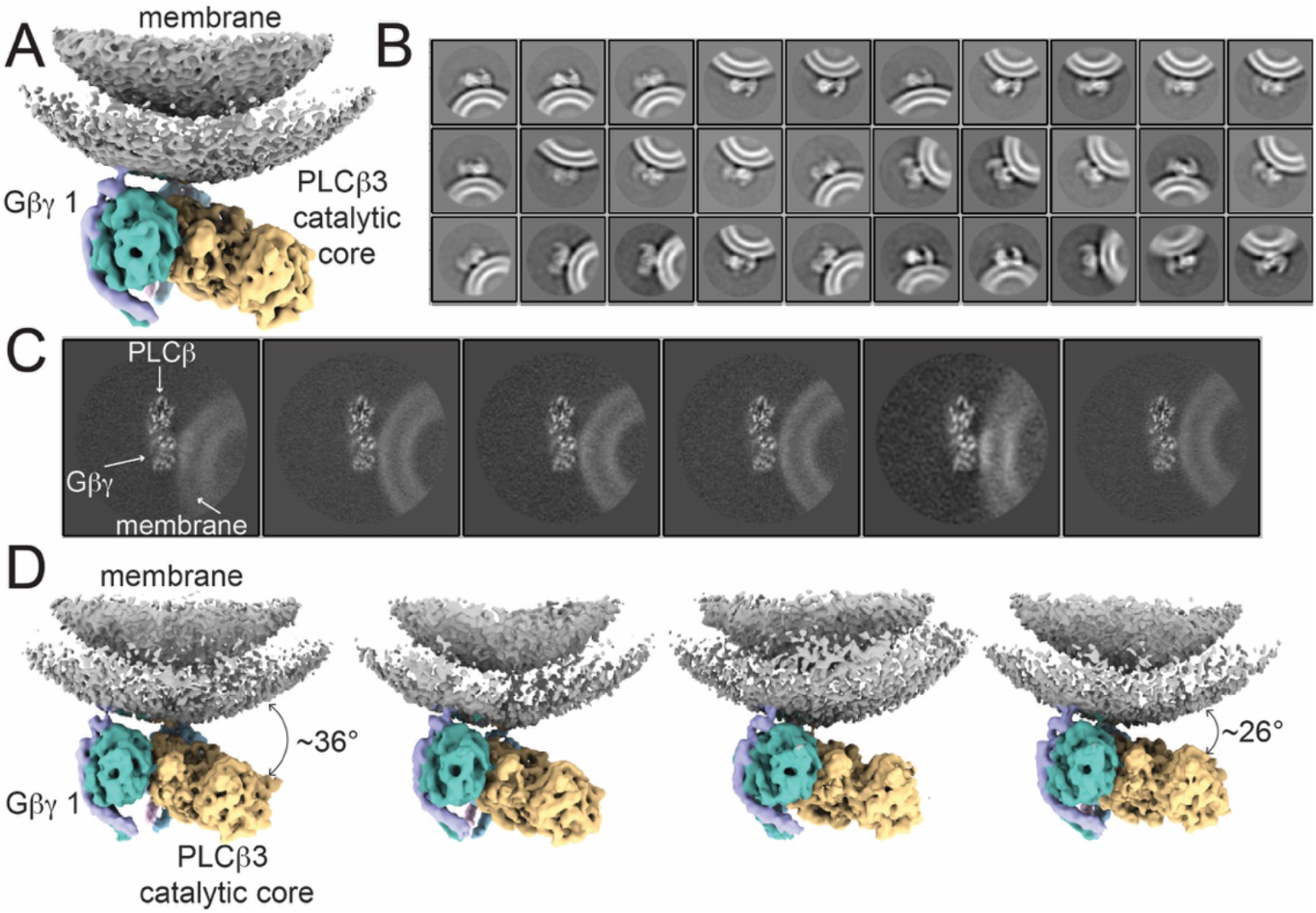
Tilting of the *PLCβ*3 ∙ *Gβγ* complex with respect to the membrane. A: Consensus unmasked refinement with density for the *PLCβ*3 ∙ *Gβγ* complex and the membrane colored by protein. The membrane is gray, *PLCβ*3 is yellow, *Gβ* 1 is dark cyan and *Gγ* 1 is light purple. B: 2D class averages of the final subset of particles determined without alignment showing side views of the complex on the membrane. Different membrane curvatures and positions of the complex with respect to the membrane are demonstrated. C: 2D projections of 3D classes of the *PLCβ*3 ∙ *Gβγ* complex on the membrane. D: 3D reconstructions of four 3D classes with different positions of the complex on the membrane arranged by degree of tilting with the most tilted on the left and least tilted on the right.

Additional 2D and 3D classification without alignment revealed heterogeneity in the position of the *PLCβ*3 ∙ *Gβγ* assembly with respect to the membrane (Figure 6B-D). 2D classes show large variation in the orientation of the catalytic core with respect to the membrane surface, with some classes showing the entire catalytic core engaged with the membrane (Figure 6B). The 2D classes also reveal differences in membrane curvature originating from differences in liposome size, which do not seem to be correlated with the degree of membrane tilting (Figure 6B). 3D classification revealed four reconstructions capturing different degrees of tilting of the catalytic core ranging from ~26° to ~36° (Figure 6D). We note that in a locally planar membrane, as opposed to a curved vesicle membrane, the active site would be nearer the membrane surface in all classes, but the variability in orientation would presumably still exist. The protein components of these reconstructions are like in the original reconstruction, with no internal conformational changes, indicating that the whole complex tilts on the membrane as a rigid body.

The lack of conformational change observed upon *Gβγ* binding and the catalytic core membrane association are consistent with our functional studies showing that activation by *Gβγ* is largely mediated by increasing membrane partitioning. Our structures suggest that the configuration of the two *Gβγ* binding sites maintains the catalytic core at the membrane and increases the probability of productive engagement with *PIP*2, potentially mediated by orientation of the catalytic core observed in our reconstructions. Taken together, our kinetic, binding, and structural studies lead us to conclude that *Gβγ* activates *PLCβ* mainly by bringing it to the membrane and orienting the catalytic core so that the active site can access the *PIP*2-containing surface (Figure 7).

**Figure 7:**
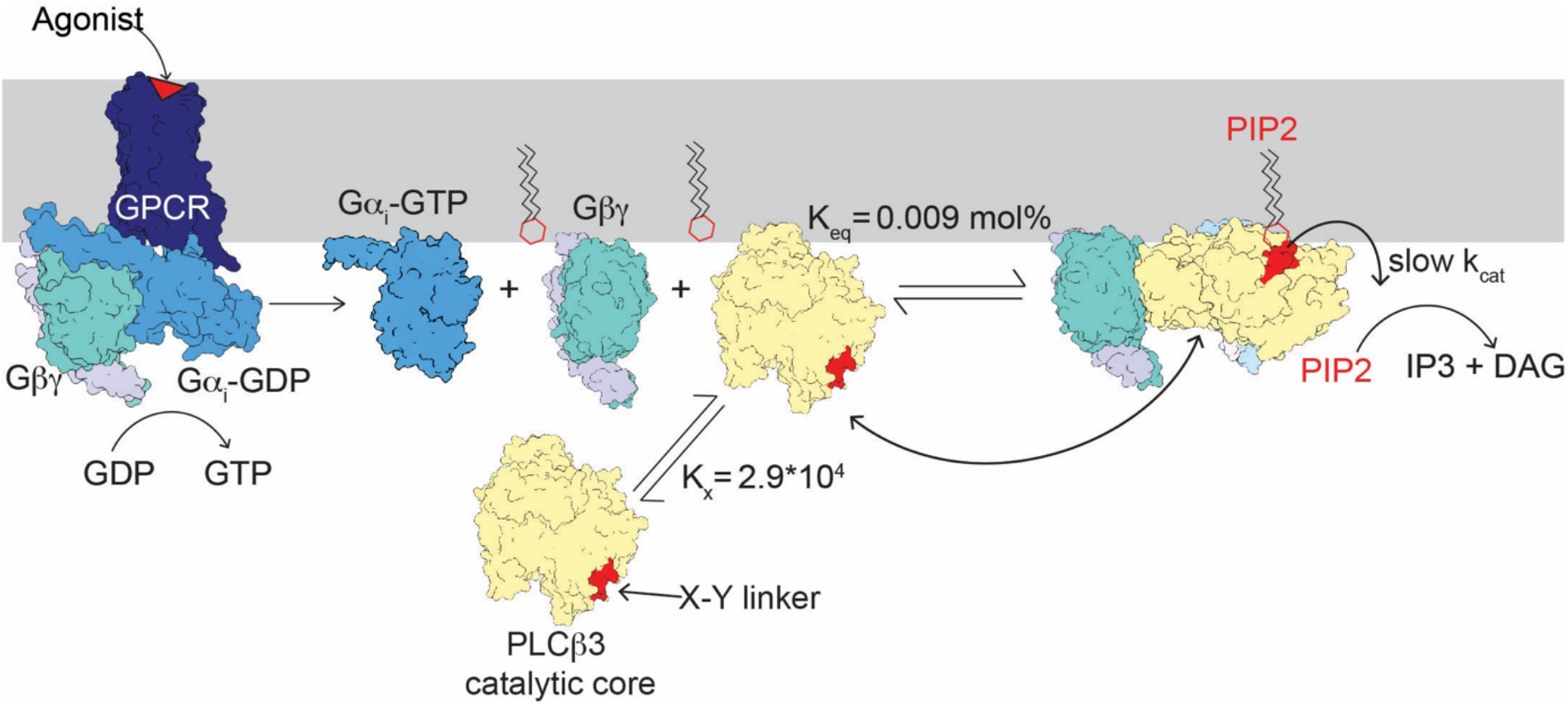
*Gβγ* activates *PLCβ* by increasing its concentration at the membrane and orienting the catalytic core to engage *PIP*2. When *PLCβ* binds the membrane, the active site is positioned away from the membrane requiring repositioning for activation, resulting in low activity in the absence in G proteins. Upon activation of a *Gα*_*i*_-coupled receptor, GTP is exchanged for GDP in the *Gα*_*i*_ subunit and free *Gβγ* is released to bind *PLCβ* which increases the concentration of *PLCβ* at the membrane and orients the active site for catalysis. The *k*_*cat*_ is limited by the X-Y linker (shown in red) which occludes the active site and is only transiently displaced from the to allow catalysis.

## Discussion

This study aims to understand how a G protein, *Gβγ*, activates the *PLCβ*3 phospholipase enzyme. We developed and applied three new technical approaches to study this process. First, because kinetic analyses of *PLCβ* enzymes historically have been limited to relatively slow radioactivity-based or semi-quantitative fluorescence assays, we have developed a new higher resolution assay using a modified, calibrated *PIP*2-dependent ion channel to provide a direct read out of membrane *PIP*2 concentration as a function of time. This assay is employed in a reconstituted system in which all components are defined with respect to composition and concentration. Second, we have used a membrane-water partition assay in, as far as we know, a novel fashion to study a surface equilibrium reaction between two proteins (*PLCβ*3 and *Gβγ*) on membranes. Third, we have determined structures of a protein complex (*PLCβ*3 and *Gβγ*) assembled on the surface of pure lipid vesicles. We also determined the structures using lipid nanodiscs, however, the lipid vesicles permitted structural analysis of the enzyme-G protein complex on lipid surfaces unperturbed by the scaffold proteins required to make nanodiscs.

We list our essential findings. (1) *PLCβ*3 catalyzes *PIP*2 hydrolysis in accordance with Michaelis-Menten enzyme kinetics. (2) *Gβγ* modifies *V*_*max*_, leaving *K*_*M*_ essentially unchanged. Under our experimental conditions, *V*_*max*_ increases ~65-fold. (3) *Gβγ* increases membrane partitioning of *PLCβ*3, an effect accountable through equilibrium complex formation between *Gβγ* and *PLCβ*3 on the membrane surface. Under our experimental conditions, partitioning increases the membrane concentration of *PLCβ*3 ~33-fold. (4) The *Gβγ*-mediated increase in *PLCβ*3 partitioning can account for most of the increase in *V*_*max*_, with a smaller, ~2-fold, effect on *k*_*cat*_. Thus, *Gβγ* regulates *PLCβ*3 mainly by concentrating it on the membrane. (5) Two *Gβγ* proteins assemble to form a complex with *PLCβ*3 on vesicle surfaces. One *Gβγ* binds to the PH domain and one EF hand of *PLCβ*3, while the other binds to the remaining EF hands. Both *Gβγ* orient their covalent lipid groups toward the membrane so that the *PLCβ*3 catalytic core is firmly anchored on the membrane surface. (6) The *PLCβ*3 ∙ *Gβγ* assembly holds the *PLCβ*3 catalytic core with its active site, as if on the end of a stylus, poised to sample the membrane surface. Assemblies on lipid vesicles reveal multiple orientations of the catalytic core with respect to the surface.

In our membrane binding analysis of the *PLCβ*3-*Gβγ* equilibrium we modeled a bimolecular reaction (1:1 stoichiometry) characterized by a single *K*_*eq*_. In our structural analysis, however, we discovered two binding sites for *Gβγ* on *PLCβ*3. Additional binding data, using multiple concentrations of *Gβγ* for example, might reveal two distinct binding constants and whether they interact with each other (i.e. behave cooperatively). Such a finding would be important because multiple binding sites could shape the *PLCβ*3 activity response to GPCR stimulation. But for purposes of the present study, the binding model treating a single site is sufficient. This is because using a single site model when two sites exist introduces an uncertainty in how *PLCβ*3 is distributed over *Gβγ*, not how much *PLCβ*3 is present in the membrane. The kinetics depend on how much *PLCβ*3 is present, and this we have measured directly with experiment.

The conclusion that *Gβγ* concentrates *PLCβ*3 on the membrane in our assay is unequivocal. To what extent do these conclusions apply to cell membranes? From Eq (8) we saw that the increase in membrane *PLCβ*3 concentration due to the fraction bound to *Gβγ* is proportional to total *Gβγ* concentration, [*G*_*tot*_]. In our assay [*G*_*tot*_] is 0.34 *mol*%, which corresponds to 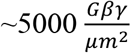. In cells we have previously estimated the concentration of *Gβγ* near GIRK2 channels in dopamine neurons during GABA_B_ receptor activation at 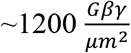 (Wang et al., 2014). Applying Eq (8), this would produce an ~9-fold increase in the membrane concentration of *PLCβ*3. This is an estimate with certain unknowns, especially the cytoplasmic concentration of *PLCβ*3 ([*PLCβ*3_*w*_]), but the result suggests that the conditions of our in-vitro assay are applicable to cell membranes. Moreover, both *Gβγ* and *Gα*_*q*_ have been shown to increase membrane association of *PLCβs* in cells, consistent with our results (Gutman et al., 2010).

We note that our demonstration that *Gβγ* increases membrane association of *PLCβ*3 directly contradicts many previous biochemical studies and the consensus in the field that G proteins do not increase the local concentration of *PLCβs* in the membrane (Jenco et al., 1997; Kadamur and Ross, 2013; Lyon and Tesmer, 2013; Romoser et al., 1996; Runnels et al., 1996). We suspect that the use of detergent solubilized *Gβγ* in past studies may have interfered with the control of its concentration on the membrane (Jenco et al., 1997; Romoser et al., 1996) (Runnels et al., 1996).

While our results and mechanism contradict the notion that G proteins do not concentrate *PLCβs* on the membrane, they are consistent with many previous observations, some we list here. As stated above, studies with cells have led to the conclusion that *Gβγ* and *Gα*_*q*_ increase membrane association of *PLCβs* (Gutman et al., 2010). The lipid anchor is required for activation of *PLCβs* by the small GPTases and Gβγ, and G proteins do not activate *PLCβs* in the absence of a membrane environment (Charpentier et al., 2014; Gutman et al., 2010; Hicks et al., 2008; Illenberger et al., 1998; Illenberger et al., 2003; Jezyk et al., 2006; Katz et al., 1992).The binding of Rac1 or *Gα*_*q*_ do not induce conformational changes around the active site, suggesting that activation is not mediated by obvious allosteric changes (Jezyk et al., 2006; Lyon et al., 2013; Waldo et al., 2010). Likewise, we observe no change in the *PLCβ*3 active site conformation when *Gβγ* is bound, only that *Gβγ* recruits *PLCβ*3 to the membrane and orients its active site.

Several properties of the *Gβγ* binding sites on *PLCβ*3 offer explanations of past observations. First, it has been shown that *Gβγ* and *Gα*_*q*_ can activate *PLCβ*3 simultaneously and additively (Carroll et al., 1995; Okajima et al., 1989; Philip et al., 2010; Rebres et al., 2011; Roach et al., 2008; Shah et al., 2009; Ware et al., 1987) and a crystal structure showed how *Gα*_*q*_ binds to *PLCβs* (Waldo et al., 2010)}(Lyon et al., 2013). We find here that the *Gβγ* sites do not occlude the *Gα*_*q*_ binding site, and therefore both G proteins can in principle bind to *PLCβ*3 simultaneously. Given the location of the *Gα*_*q*_ site relative to the *Gβγ* sites and the active site, we suspect that when three G-proteins are bound (2 *Gβγ* and 1 *Gα*_*q*_), the active site will be brought into closer proximity to the membrane surface than when only the *Gβγ* are bound, explaining enhanced catalysis. Second, several amino acids on *Gβγ* that contact *PLCβ*3 in the structure were previously shown to play a role in binding to *Gα*, *PLCβ* and other effectors (Figure 5C-D) (Ford et al., 1998). Third, the PH domain was shown to play a role in *Gβγ* binding and activation, however, based on our structures, *Gβγ* binding does not require or induce rearrangement of the catalytic core as was previously proposed (Drin et al., 2006; Kadamur and Ross, 2016). Fourth, Rac1 was also shown to bind to the PH domain of *PLCβ*2 (Figure S6D), and Rac1-activated *PLCβ* was shown to be additionally activated by *Gβγ*, leading to a proposal that the two binding sites did not overlap (Illenberger et al., 2003; Jezyk et al., 2006). Our structures show that Rac1 and *Gβγ* do indeed share an interface within the PH domain (Figure S6D), however, the second *Gβγ* binding site can explain the additive activation (Illenberger et al., 2003; Jezyk et al., 2006).

An intriguing aspect of *PLCβ* enzymes is that all wild type structures show that the active site is occluded by the inhibitory X-Y linker. This includes complexes with *Gα*_*q*_ Rac1, and now *Gβγ* (Jezyk et al., 2006; Lyon et al., 2013; Lyon et al., 2014b; Waldo et al., 2010). It has been proposed that lipids are required to remove the X-Y linker to achieve catalysis (Charpentier et al., 2014; Lyon et al., 2014b; Lyon and Tesmer, 2013). This must be true because unless the linker is displaced, catalysis cannot occur. From our data, we put forth a proposal that is consistent with this autoinhibition idea but lends new insight into this system. Our quantitative kinetic analysis shows that in a perfectly intact lipid membrane environment without detergent, that *k*_*cat*_ for *PLCβ*3, 1.7 *sec*^−1^in the absence and 3.2 *sec*^−1^ in the presence of *Gβγ* (Figure 3C), is extremely small. We propose that *PLCβs* have evolved to have a small *k*_*cat*_ produced by this autoinhibitory element. This ensures that in the absence of GPCR stimulation, the baseline partitioning of *PLCβ* enzyme from the cytoplasm to the membrane, determined by *K*_*x*_ and the cytoplasmic concentration of *PLCβ*, produces very little *PIP*2 hydrolysis. Only upon GPCR stimulation, when a large quantity of *PLCβ* partitions into the membrane, determined by *K*_*eq*_ and the *Gβγ* concentration generated by GPCR stimulation (and a similar contribution by *Gα*_*q*_), is there enough *PLCβ* enzyme in the membrane, even though *k*_*cat*_ remains low, to catalyze *PIP*2 hydrolysis. In other words, a small *k*_*cat*_ combined with an ability to enact large changes in membrane enzyme concentration upon GPCR stimulation permits a strong signal when the system is stimulated and a minimal baseline when it is not.

## Supporting information

primary SI

## Acknowledgements

We thank Chen Zhao for developing and characterizing the ALFA nanobody mediated tethering of *Gβγ* to GIRK and for insightful discussions. We thank Venkata S. Mandala for assistance with protein reconstitution and NMR experiments. We thank Christoph A. Haselwandter for insightful discussion and comments on the manuscript. We thank Yi Chun Hsiung for assistance with tissue culture. We thank members of the MacKinnon lab, Jue Chen and members of her lab for helpful discussions. This work was supported by NIHF32GM142137 to MEF. RM is an investigator in the Howard Hughes Medical Institute.

We thank Rui Yan and Zhiheng Yu at the HHMI Janelia Cryo-EM Facility for help in microscope operation and data collection.

We thank Mark Ebrahim, Johanna Sotiris, and Honkit Ng at the Evelyn Gruss Lipper Cryo-EM Resource Center of Rockefeller University for assistance with cryo-EM data collection.

Some of this work was performed at the Simons Electron Microscopy Center and National Resource for Automated Molecular Microscopy located at the New York Structural Biology Center, supported by grants from the Simons Foundation (SF349247), NYSTAR, and the NIH National Institute of General Medical Sciences (GM103310) with additional support from Agouron Institute (F00316) and NIH (OD019994).

## Author contributions

R.M. and M.E.F. designed studies; M.E.F. performed experiments; R.M. and M.E.F. analyzed and interpreted data; R.M. and M.E.F. wrote the paper.

## Star Methods

### Resource Availability

#### Lead contact

Further information and requests for resources and reagents should be directed to and will be fulfilled by the lead contact, Dr. Roderick MacKinnon

#### Materials availability

This study did not generate new unique reagents.

#### Data and code availability

Cryo-EM maps and atomic models for all structures described in this work have been deposited to the Electron Microscopy Data Bank (EMDB) and the Protein Data Bank (PDB), respectively. Accession codes are as follows: *PLCβ*3 in solution-8EMV and EMD-28266, *PLCβ*3 in complex with *Gβγ* on vesicles-8EMW and EMD-28267, and *PLCβ*3 in complex with *Gβγ* on nanodiscs-8EMX and EMD-28268.

This work does not report original code.

Any additional information required to reanalyze the data reported in this paper is available from the lead contact upon request.

### Experimental Models and Subject Details

Sf9 cells (Cellosaurus ID: CVCL_0549, female) were cultured in Sf-900 II SFM medium supplemented with 100 U/mL penicillin and 100 U/mL streptomycin at 27 °C. High Five cells (Cellosaurus ID: CVCL_C190, sex unspecified) were cultured in Express Five SFM media supplemented with 10 mM L-glutamine, 100 U/mL penicillin and 100 U/mL streptomycin at 27 °C. HEK293S GnTl-cells (Cellosaurus ID: CVCL_A785, female) were cultured in Freestyle 293 medium supplemented with 2% fetal bovine serum (FBS), 100 U/mL penicillin and 100 U/mL streptomycin at 37 °C.

### Method Details

#### Protein Expression and Purification

The human *PLCβ*3 gene was provided by Dr. Sondek (Charpentier et al., 2014) and contained residues 10 to 1234. It was cloned into the pFastBac vector downstream of GFP and transformed into DH10bac cells to produce bacmid DNA, which was used to transfect Sf9 cells to produce baculovirus. High Five insect cells at 2×10^6^ cells/mL were infected with 15-25 *mL* of P3 virus per liter of culture and harvested 36-48 hours after infection by centrifugation at 3,500 x *g* for 15 minutes. Pellets were flash frozen and stored at −80°C until use. Purification was carried out at 4°C. Cells were resuspended in 100 *mL* of buffer containing 50 *mM* HEPES pH 8.0, 50 *mM* NaCl, 10 *mM* 2-mercaptoethanol, 5% glycerol (v/v), 0.1 *mM* EDTA, 0.1 *mM* EGTA, DNase and protease inhibitors (12.5 *μg*/*mL* leupeptin, 12.5 *μg*/*mL* pepstatin A, 625 *μg*/*mL* AEBSF, 1 *mM* Benzamadine, 100 *μg*/*mL* Trypisin inhibitor, 1x aprotinin, and 1 *mM* PMSF) and lysed by brief sonication. Lysate was clarified by centrifugation at 39,000 x *g* for 45 minutes and bound to GFP nanobody-coupled Sepharose resin (prepared in-house) for one hour. The resin was washed in batch once with 10 column volumes of buffer containing 20 *mM* HEPES pH 8.0, 400 *mM* NaCl, 10 *mM* 2-mercaptoethanol, 2% glycerol (v/v), 0.1 *mM* EDTA, 0.1 *mM* EGTA, and protease inhibitors (625 *μg*/*mL* AEBSF, 1 *mM* Benzamadine, 100 *μg*/*mL* Trypisin inhibitor, and 1x aprotinin) then loaded onto a column and washed with an additional 10 column volumes by gravity flow. Protein was eluted by cleavage with 3C PreScission protease (made in house) for two hours, concentrated to ~10 mg/mL using a 15-mL Amicon concentrator with 100-kDa molecular weight cutoff, and further purified by size exclusion chromatography using a Superdex 200 10/300 increase column in buffer containing 20 mM HEPES pH 8.0, 100 mM NaCl, 5 mM Dithiothreitol (DTT), 2% glycerol (v/v), 0.1 mM EDTA, 0.1 mM EGTA, and protease inhibitors (12.5μg/mL leupeptin, 12.5μg/mL pepstatin A, 625μg/mL AEBSF, 1 mM Benzamadine, 100 μg/mL Trypisin inhibitor, 1x aprotinin, and 1 mM PMSF). Fractions with *PLCβ*3 were pooled, flash frozen, and stored at −80°C for later use. For preparation of cryo-EM grids, protein was used directly after size exclusion.

To purify protein for non-specific cysteine labeling, the 10 *mM* 2-mercaptoethanol in all buffers was replaced with 2 *mM* tris(2-carboxyethyl)phosphine (TCEP) and the protein-loaded resin was washed with buffer containing 20 *mM* HEPES pH 7.4, 400 *mM* NaCl, 2 *mM* TCEP, 2% glycerol (v/v), 0.1 *mM* EDTA, 0.1 *mM* EGTA, and protease inhibitors (625 *μg*/*mL* AEBSF, 1 *mM* Benzamadine). Following elution, maleimide LD655 (Zheng et al., 2016) was added in 5-fold molar excess and incubated overnight protected from light. Labeled protein was concentrated ~10 *mg*/*mL* using a 15-mL Amicon concentrator with 100-kDa molecular weight cutoff, and further purified by size exclusion chromatography using a superdex 200 10/300 increase column in buffer containing 20 mM HEPES pH 8.0, 100 mM NaCl, 5 mM Dithiothreitol (DTT), 2% glycerol (v/v), 0.1 *mM* EDTA, 0.1 *mM* EGTA, and protease inhibitors 12.5 *μg*/*mL* leupeptin, 12.5 *μg*/*mL* pepstatin A, 625 *μg*/*mL* AEBSF, 1 *mM* Benzamadine, 100 *μg*/*mL* Trypisin inhibitor, and 1x aprotinin). Fractions with labeled *PLCβ*3 were pooled, flash frozen, and stored at −80°C for later use. Labeling efficiency was consistently 60-70%.

For lipidated *Gβγ*, untagged human *Gβ*1 was co-expressed with bovine *Gγ*2 with an N-terminal His-YFP tag in High Five insect cells using 12 and 8 *mL* of P3 baculovirus respectively at 2×10^6^ cells/mL for 36-48 hours. Cells were harvested by centrifugation at 3,500 x *g* for 15 minutes and pellets were flash frozen and stored at −80°C until use. Purification was carried out at 4°C. ~45 *mL* of cells were resuspended in 200 *mL* of buffer containing 25 *mM* Tris-HCl pH 8.0, 125 *mM* NaCl, 5 *mM* EGTA, 5 *mM* DTT, DNase and protease inhibitors (12.5 *μg*/*mL* leupeptin, 12.5 *μg*/*mL* pepstatin A, 625 *μg*/*mL* AEBSF, 1 *mM* Benzamadine, 100 *μg*/*mL* Trypisin inhibitor, 1x aprotinin, and 1 *mM* PMSF) and broken by manual homogenization. Membranes were separated by centrifuging at 39,000 x *g* for 30 minutes, resuspended in buffer containing 25 *mM* Tris-HCl pH 8.0, 125 *mM* NaCl, DNase and protease inhibitors (12.5 *μg*/*mL* leupeptin, 12.5 *μg*/*mL* pepstatin A, 625 *μg*/*mL* AEBSF, 1 *mM* Benzamadine, 100 *μg*/*mL* Trypisin inhibitor, 1x aprotinin, and 1 *mM* PMSF) and manually homogenized again. Proteins were extracted using 1% sodium cholate added from a 10% stock by mixing for 1.5 hours. Lysate was clarified by centrifuging at 39,000 x *g* for 30 minutes and bound in batch to 10 *mL* TALON resin equilibrated with buffer containing 25 *mM* Tris-HCl pH 8.0, 125 *mM* NaCl, and 1% sodium cholate for one hour. The resin was washed in batch with 100 *mL* of equilibration buffer then loaded onto a column and washed by gravity flow with 10 column volumes of high salt buffer (25 *mM* Tris-HCl pH 8.0, 500 *mM* NaCl, and 1% sodium cholate) and 10 *mM* imidazole buffer (25 *mM* Tris-HCl pH 8.0, 125 *mM* NaCl, 1% sodium cholate, and 10 *mM* imidazole). Protein was eluted with ~20 *mL* buffer containing 25 *mM* Tris-HCl pH 8.0, 125 *mM* NaCl, 1% sodium cholate, and 200 *mM* imidazole and concentrated to ~2 *mL* using a 15-*mL* Amicon concentrator with 30-kDa molecular weight cutoff. The final 2 *mL* of protein was diluted to ~20 *mL* using buffer without imidazole and incubated with 3C PreScission protease (prepared in-house) overnight at 4°C remove the His-YPF. The cleaved protein was passed over a column of 10 *mL* TALON resin equilibrated with 25 *mM* Tris-HCl pH 8.0, 125 *mM* NaCl, 1% sodium cholate, and 20 *mM* imidazole three times to remove the free His-YFP, and concentrated to 1 *mL* using a 15-mL Amicon concentrator with 30-kDa molecular weight cutoff. *Gβγ* was purified further via size exclusion chromatography using a Superdex 200 10/300 increase column in buffer containing 25 *mM* Tris-HCl pH 8.0, 125 *mM* NaCl, 1% sodium cholate, and 5 *mM* DTT. Fractions containing *Gβγ* were pooled and concentrated to 5-10 *mg*/*mL* using 4-mL Amicon concentrator with 30-kDa molecular weight cutoff and immediately used for reconstitution.

For nanodisc reconstitution, the YFP was maintained to facilitate purification of protein-containing nanodiscs. Following elution from TALON resin, protein was concentrated to 1 *mL* using a 15-mL Amicon concentrator with 30-kDa molecular weight cutoff and further purified by size exclusion chromatography using a Superdex 200 10/300 increase column in buffer containing 25 *mM* Tris-HCl pH 8.0, 125 *mM* NaCl, 1% sodium cholate, and 5 *mM* DTT. Fractions containing *Gβγ* −YFP were pooled and concentrated to ~10 *mg*/*mL* using 4-mL Amicon concentrator with 30-kDa molecular weight cutoff and immediately used for reconstitution.

For soluble *Gβγ*, untagged human *Gβ*1 was co-expressed with human *Gγ*2 C68S with an N-terminal His-YFP tag in High Five insect cells using 12 and 8 mL of P3 baculovirus respectively at 2×10^6^ cells/mL for 36-48 hours. Cells were harvested by centrifugation at 3,500 x *g* for 15 minutes and pellets were flash frozen and stored at −80°C until use. Purification was carried out at 4°C. ~45 *mL* of cells were resuspended in 200 *mL* of buffer containing 25 *mM* Tris-HCl pH 8.0, 125 *mM* NaCl, DNase and protease inhibitors (12.5 *μg*/*mL* leupeptin, 12.5 *μg*/*mL* pepstatin A, 625 *μg*/*mL* AEBSF, 1 *mM* Benzamadine, 100 *μg*/*mL* Trypisin inhibitor, 1x aprotinin, and 1 *mM* PMSF) and broken brief sonication. Lysate was clarified by centrifuging at 39,000 x *g* for 45 minutes and bound in batch to 10 *mL* TALON resin equilibrated with buffer containing 25 *mM* Tris-HCl pH 8.0, and 125 *mM* NaCl for one hour. The resin was washed in batch with 100 *mL* of equilibration buffer then loaded onto a column and washed by gravity flow with 10 CV of high salt buffer (25 *mM* Tris-HCl pH 8.0, and 500 *mM* NaCl) and 10 *mM* imidazole buffer (25 *mM* Tris-HCl pH 8.0, 125 *mM* NaCl, and 10 *mM* imidazole). Protein was eluted with ~20 *mL* buffer containing 25 *mM* Tris-HCl pH 8.0, 125 *mM* NaCl, and 200 *mM* imidazole and concentrated to ~2 *mL* using a 15-mL Amicon concentrator with 30-kDa molecular weight cut off. The final 2 *mL* of protein was diluted to ~20 *mL* using buffer without imidazole and incubated with 3C PreScission protease (prepared in-house) overnight at 4°C to remove the His-YPF. The cleaved protein was passed over a column of 10 *mL* TALON resin equilibrated with 25 *mM* Tris-HCl pH 8.0, 125 *mM* NaCl, and 20 *mM* Imidazole three times to remove the free His-YFP, and concentrated to 1 *mL* using a 15-mL Amicon concentrator with 30-kDa molecular weight cutoff. *Gβγ* was purified further using size exclusion chromatography using a superdex 200 10/300 increase column in buffer containing 25 *mM* Tris-HCl pH 8.0, 125 *mM* NaCl, and 5 *mM* DTT. Fractions containing *Gβγ* were pooled, flash frozen, and stored at - 80°C until use.

To attempt to form a complex between *PLCβ*3 and soluble *Gβγ*, proteins were mixed at a 2:1 molar ratio of *Gβγ* : *PLCβ*3, incubated on ice for one hour and run on size exclusion chromatography using a Superdex 200 10/300 increase column in buffer containing 20 *mM* HEPES pH 8.0, 100 *mM* NaCl, 5 *mM* Dithiothreitol (DTT), 2% glycerol (v/v), 0.1 *mM* EDTA, 0.1 *mM* EGTA, and protease inhibitors (112.5 *μg*/*mL* leupeptin, 12.5 *μg*/*mL* pepstatin A, 625 *μg*/*mL* AEBSF, 1 *mM* Benzamadine, 100 *μg*/*mL* Trypisin inhibitor, 1x aprotinin, and 1 *mM* PMSF). Peaks were analyzed by SDS-PAGE and showed no-comigration (Figure S3E).

For ALFA-nanobody tagged *Gβγ*, untagged human *Gβ*1 was co-expressed with human *Gγ*2 with the ALFA nanobody on the N-terminus and an N-terminal His-YFP tag in High Five insect cells using 12 and 8 mL of P3 baculovirus respectively at 2×10^6^ cells/mL for 36-48 hours. Cells were harvested by centrifugation at 3,500 x *g* for 15 minutes and pellets were flash frozen and stored at −80°C until use. Purification was carried out at 4°C. ~45 *mL* of cells were resuspended in 200 *mL* of buffer containing 25 *mM* Tris-HCl pH 8.0, 125 *mM* NaCl, 5 *mM* DTT, DNase and protease inhibitors (12.5 *μg*/*mL* leupeptin, 12.5 *μg*/*mL* pepstatin A, 625 *μg*/*mL* AEBSF, 1 *mM* Benzamadine, 100 *μg*/*mL* Trypisin inhibitor, 1x aprotinin, and 1 *mM* PMSF) and broken by brief sonication. Lysate was clarified by centrifugation at 39,000 x *g* for 45 minutes and bound to GFP nanobody-coupled Sepharose resin (prepared in house) for one hour. The resin was washed in batch one time with 10 column volumes of buffer containing 25 *mM* Tris-HCl pH 8.0, 125 *mM* and NaCl, then loaded into a column and washed with an additional 10 column volumes by gravity flow. Protein was eluted by cleavage with 3C PreScission protease (prepared in-house) for two hours, concentrated to 1 mL using a 15-mL Amicon concentrator with 30-kDa molecular weight cutoff, and further purified by size exclusion chromatography using a Superdex 200 10/300 increase column in buffer containing 25 *mM* Tris-HCl pH 8.0, 125 *mM* NaCl, and 5 *mM* DTT. Fractions with nanobody-tagged *Gβγ* were pooled, flash frozen, and stored at −80°C for later use.

For ALFA-tagged GIRK, full-length mouse GIRK2 with a C-terminal ALFA peptide tag upstream of a GFP tag was expressed using HEK293S GnTI^−^ cells at a density of ~3.5*10^6^ cells/mL infected with 10% (v/v) P3 virus. 10 *mM* sodium butyrate was added 12 hours after infection and the temperature was reduced to 30°C for 48 hours. Cells were harvested by centrifugation at 3,500 x *g* for 15 minutes and pellets were flash frozen and stored at −80°C until use. Purification was carried out at 4°C. ~30 *mL* of cells were resuspended in buffer containing 25 *mL* Tris pH 7.5, 150 *mM* KCl, 2 *mM* DTT, DNase and protease inhibitors (12.5 *μg*/*mL* leupeptin, 12.5 *μg*/*mL* pepstatin A, 625 *μg*/*mL* AEBSF, 1 *mM* Benzamadine, 100 *μg*/*mL* Trypisin inhibitor, 1x aprotinin, and 1 *mM* PMSF) and lysed by manual homogenization. Membranes were separated by centrifugation at 39,000 x *g* for 30 minutes, resuspended in buffer containing 20 *mM* Tris pH 7.5, 150 *mM* KCl, 2 *mM* DDT, and manually homogenized. Channels were extracted with 1.5% DDM/0.3% CHS for 1.5 hours and lysate was clarified by centrifugation 39,000 x *g* for 30 minutes. Lysate was bound to GFP nanobody-coupled Sepharose resin (prepared in house) for one hour, washed in batch one time with 10 column volumes of buffer containing 20 *mM* Tris pH 7.5, 150 *mM* KCl, 2 *mM* DDT, and 0.05%/0.01% DDM/CHS, then loaded into a column and washed with an additional 10 column volumes by gravity flow. Protein was eluted by cleavage with 3C PreScission protease (prepared in-house) for two hours, concentrated to ~10 *mg*/*mL* using a 15-mL Amicon concentrator with 100-kDa molecular weight cutoff, and further purified by size exclusion chromatography using a Superose 6 10/300 increase column in buffer containing 20 *mM* Tris pH 7.5, 150 *mM* KCl, 10 *mM* DDT, and 0.025%/0.005% DDM/CHS. Fractions with GIRK2 were pooled, concentrated to 2 *mg*/*mL* and used for reconstitution immediately.

#### Protein Reconstitution

GIRK and lipidated *Gβγ* were reconstituted for bilayer experiments using 3:1 ratio of 1-palmitoyl-2-oleoyl-sn-glycero-3-phosphoethanolamine (POPE): 1-palmitoyl-2-oleoyl-sn-glycero-3-phospho-(1’-rac-glycerol) (POPG) lipids. Lipids in chloroform were mixed and dried under a stream of argon, washed with pentane, dried under a stream of argon and incubated under vacuum overnight. Lipids were resuspended at 20 *mg*/*mL* in buffer containing 10 *mM* K_2_HPO_4_, 450 *mM* KCl, and 10 *mM* DTT and sonicated to clarity. 1% DM was added to the lipids and the mixture was sonicated extensively. GIRK was added to the mixture at a protein to lipid ratio of 1:10 (wt/wt), or *Gβγ* was added at 1:5 (wt/wt), the lipid concentration was diluted to 10 *mg*/*mL* and incubated at 4°C for one hour. Detergent was removed at 4°C using dialysis in a tubing with a 50 kDa cutoff for GIRK or a 10 kDa cutoff for *Gβγ* in buffer containing 10 *mM* K_2_HPO_4_ pH 7.4, 450 *mM* KCl, and 10 *mM* DTT, and biobeads. Dialysis buffer was changed every 12 hours for three changes with fresh DTT added at each change. After 3 changes, liposomes were incubated with ~50% volume of biobeads for five hours at 4°C, flash frozen, and stored at −80°C until use.

For structural studies using liposomes, *Gβγ* was reconstituted using a 2:1:1 mixture of 1,2-dioleoyl-sn-glycero-3-phosphoethanolamine (DOPE): 1-palmitoyl-2-oleoyl-glycero-3-phosphocholine (POPC): 1-palmitoyl-2-oleoyl-sn-glycero-3-phospho-L-serine (POPS). Lipids in chloroform were mixed and dried under a stream of argon, washed with pentane, dried under a stream of argon, and incubated under vacuum overnight. Lipids were resuspended at 25 *mM* in buffer containing 25 *mM* HEPES pH 7.4, 150 *mM* KCl and 5 *mM* DTT and sonicated to clarity. 40 *mM* sodium cholate was added and the mixture was sonicated briefly. *Gβγ* was added at a protein to lipid ratio of 1:15 (wt/wt) and the lipid concentration was reduced to 20 *mM* maintaining 40 *mM* sodium cholate and incubated at 4°C for one hour. Detergent was removed using four exchanges of 200 *mg*/*mL* biobeads washed with reconstitution buffer after two hours, 12 hours, two hours, and two hours at 4°C. Liposomes were used immediately following reconstitution to prepare grids. Reconstitution for partition experiments was carried out in the same way but 0.1 *mole*% 1,2-dioleoyl-sn-glycero-3-phosphoethanolamine-N-(lissamine rhodamine B sulfonyl) (18:1 Liss Rhod PE) was included, the final lipid concentration was diluted to 12.5 *mM*, and *Gβγ* was added at a protein to lipid ratio of 1:5 (wt/wt). Liposomes for partitioning studies were flash frozen and stored at −80°C until use.

For nanodiscs, 2DOPE:1POPC:1POPS lipids were used. Lipids in chloroform were mixed and dried under a stream of argon, washed with pentane, dried under a stream of argon, and incubated under vacuum overnight. Lipids were resuspended at 20 *mM* in buffer containing 25 *mM* HEPES pH 7.4, 150 *mM* KCl, 5 *mM* DTT, and 40 *mM* sodium cholate and sonicated to clarity. MSP2N2 scaffold (Grinkova et al., 2010), *Gβγ*-YFP, and lipids, were mixed at a molar ratio of 1:0.75:220 in buffer containing 25 *mM* HEPES pH 7.4, 150 *mM* KCl, and 5 *mM* DTT and incubated for one hour at 4°C. Detergent was removed using two incubations with 200 *mg*/*mL* biobeads washed with reconstitution buffer changed after two hours and 12 hours. Harvested nanodiscs were diluted to 3 *mL* with buffer containing 5 *mM* HEPES pH 7.4, 150 *mM* KCl, and 5 *mM* DTT and bound in batch to GFP nanobody-coupled Sepharose resin (prepared in house) for one hour. Resin was loaded onto a column and washed with 10 column volumes of 25 *mM* HEPES pH 7.4, 150 *mM* KCl, and 5 *mM* DTT buffer by gravity flow and eluted by cleavage with 3C PreScission protease (prepared in-house) for two hours. Nanodiscs were concentrated to 1 *mL* using a 4-mL Amicon concentrator with 100-kDa molecular weight cutoff, and further purified by size exclusion chromatography using a Superose 6 10/300 increase column in buffer containing 25 *mM* HEPES pH 7.4, 150 *mM* KCl, and 5 *mM* DTT.

Reconstitution of *Gβγ* was confirmed using SDS PAGE, nanodiscs were concentrated to 5 *mg*/*mL* using a 0.5-mL Amicon concentrator with 100-kDa molecular weight cutoff and used immediately for grid preparation.

#### Bilayer experiments and analysis

A lipid mixture of 2DOPE:1POPC:1POPS with various concentrations of L-α-phosphatidylinositol-4,5-bisphosphate (Brain, Porcine) (*PIP*2) was used. Lipids in chloroform were dried under a stream of argon and resuspended at 20 *mg*/*mL* in decane. A vertical bilayer configuration was used for all experiments. Two chamber cups (3 *mL* each) were connected by a 100 *μm* thick piece of Fluorinated ethylene propylene copolymer with a ~250 μm hole, which was used to paint a bilayer with the lipid-decane mixture. Buffer containing 25 *mM* HEPES pH 7.4, 150 *mM* KCl, 30 *mM* NaCl, 2 *mM* MgCl_2_, and 100 *μM* CaCl_2_ was used in both chambers. The reference electrode and the ground electrode were connected to the trans and cis chambers respectively via agarose salt bridges. A magnetic stirbar was included in the cis chamber to facilitate mixing. Voltage across the lipid bilayer was controlled with an Axopatch 200Bamplifier in whole-cell mode. The analog current signal was lowpass filtered at 1 kHz (Bessel) and digitized at 10 kHz with a Digidata 1440A digitizer. Digitized data were recorded with software pClamp (MolecularDevices). 30 *nM* Nanobody-tagged *Gβγ* was added at the beginning of each experiment. All reagents (GIRK-containing vesicles, *PLCβ*3, etc) were added to the cis chamber under continuous mixing. Frozen GIRK-containing liposomes were thawed on ice and KCl solution was added to a final concentration of 1 M. Before fusing liposomes, they were sonicated briefly at room temperature.

For the *PIP*2 titration experiments, the starting concentration of *PIP*2 in the bilayer was varied from 0.1 *mol*% to 4 *mol*%. For each experiment, GIRK-containing vesicles were fused and current at +80 mV was measured. While recording, 32 *μM* C8PIP2 was added to maximally activate the channels. The starting current was normalized to the maximal current and reported as I/Imax. Each starting *PIP*2 concentration was repeated at 3-6 times. For *PLCβ*3 experiments, 1 *mol*% *PIP*2 was used. After fusing vesicles with GIRK, baseline current at +80 mV was measured for 3-5 minutes and then 29 *nM* (1x) or 58 *nM* (2x) *PLCβ*3 was added to the cis chamber while recording. After the current decay, a voltage family was measured to ensure integrity of the bilayer and then 32 μM *C*8*PIP*2 was added to recover channel activity. Any experiment where the current did not recover was discarded. For experiments with soluble *Gβγ*, 1 *μM* was added before the addition of *PLCβ*3. Analysis was carried out in Clampfit. Current decays were filtered at 10-20 hz, reduced by a factor of 10 and exported to qtiplot. Initially, the leak was subtracted and the decays were normalized to the starting *PIP*2 concentration and converted to *PIP*2 decays using Equation 1. For Michaelis-Menten analysis, current decays were normalized to the starting *PIP*2 concentration and directly fit to Equation 4 to determine *K*_*M*_ and *V*_*max*_.

#### NMR experiments to measure lipid concentration

A small volume of liposomes from reconstitutions (10-20 μL) was dissolved in ~550 uL of a mixture of deuterated methanol and chloroform (5:1 or 4:1) containing 100 μM of the standard sodium trimethylsilyl propionate (TSP). Proton spectra were measured on a Bruker 600 MHz instrument equipped with an AVANCE NEO console and a 5 mm HCN cryoprobe. Spectra were collected in 5 mm tubes at 298 K using a 30° flip angle, 16 scans, and 2.8 second acquisition time and a recycle delay of 18 μs. Spectra were processed using TopSpin 4.1.1 for line broadening, phasing, and baseline correction. The triplet lipid -CH_3_ peak at 0.875 ppm was integrated relative to the TSP peak at 0 ppm and normalized to the difference in protons (nine for TSP and 6 for the lipid CH3) (Figure S2A). The normalized peak area was used to determine the lipid concentration using the known 100 *μM* concentration of TSP.

#### PLCβ vesicle partition experiments

Reconstituted liposomes with or without *Gβγ* were subjected to 10 freeze-and-thaw cycles and extruded 21 times through a 200 nm membrane to produce LUVs. Fixed concentrations of lipids were mixed with LD655-labeled *PLCβ*3 in buffer containing 25 *mM* HEPES pH 7.4, 150 *mM* KCl, and 5 *mM* DTT, incubated for one hour, and centrifuged for one hour at 100,000 x *g* at room temperature. The supernatant was removed and the membrane pellet was resuspended in an equal volume of buffer. The input, pellet, and supernatant samples were analyzed by SDS-page and imaged using in-gel fluorescence to detect LD655-labeled *PLCβ* (Figure S2D-E). Gel bands were quantified using Biorad imagelab software. Additionally, input, supernatant, and pellet samples were solubilized in 3-5% Anapoe-C12E10 to eliminate scattering artifacts and the fluorescence signal was measured for LD655 (ext-649, em-666) and Rhodamine (ext-560, em-583) using a Tecan plate reader. The Rhodamine signal was used to estimate the fraction of lipids that were pelleted and the measurements were corrected for this as well as the loss of material using the difference between the input and output (pellet and supernatant) LD655 signal. The reported lipid concentration is 50% of the total lipid concentration added in solution because *PLCβ*3 only has access to the outer leaflet. Each lipid concentration was repeated with two different *PLCβ*3 concentrations, 100 *nM* and 300 *nM*. In the absence of *Gβγ*, values for *K*_*x*_ were determined for each experiment using Equation 5 and values for fraction of *PLCβ*3 partitioned (*F*_*p*_), were determined, plotted against lipid concentration, and fit to Equation 6 for a more robust determination of *K*_*x*_ (Figure 3D, S2A-B). In the presence of *Gβγ*, values for 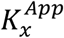 were determined for each experiment using Equation 5 and values for fraction of *PLCβ*3 partitioned (*F*_*p*_), were determined, plotted against lipid concentration, and fit to Equation 7 for *K*_*eq*_ (Figure 3D, S2A-B). Values from the gels and the solution fluorescence measurements were consistent so the values from the solution measurements are reported. Individual values for *K*_*x*_ and 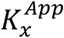 do not vary with *PLCβ*3 or lipid concentrations, as expected (White et al., 1998). Further, values of *F*_*p*_, *K*_*x*_, and 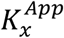 were consistent across multiple reconstitutions and preparations (Figure S2B-D).

#### Cryo-EM sample preparation and data collection

For *PLCβ*3 in solution without liposomes, glycerol was omitted from the size exclusion buffer and protein was concentrated to 4.8 *mg*/*mL* using a 4-mL Amicon concentrator with 100-kDa molecular weight cutoff. Protein was supplemented with 3 mM of Fluorinated Fos-Choline-8 ~5 minutes before grid preparation. Quantifoil R1.2/1.3 400 mesh holey carbon Au grids were glow discharged for 20s, and 3.5 μL of sample was applied. After 20s incubation at 16°C and 100% humidity, grids were blotted for 2.5s with a blot force of 0, and plunge frozen in liquid ethane using a FEI Vitrobot Mark IV. For data acquisition, grids were loaded onto a 300-kV Titan Krios transmission electron microscope with a Gatan K3 Summit direct electron detector and a GIF quantum energy filter with a slit width of 20 eV and 3,527 movies were collected in superresolution mode with a pixel size of 0.54 Å and a defocus range of 1 to 2.5 *μm* using SerialEM (Mastronarde, 2005). The movies were recorded with 40 frames, a 2 second total exposure time (0.05s/frame) and a dose rate of 25 e^−^/pix/s which gave a cumulative dose of 42.87 e^−^/Å^2^ (1.07 e^−^/Å^2^/frame).

For *PLCβ*3 and reconstituted liposomes without *Gβγ*, *PLCβ*3 and liposomes were mixed at final concentrations of 0.5 *mg*/*mL* and 17.5 *mM* respectively and incubated at room temperature for one hour. The sample was supplemented with 3 *mM* of Fluorinated Fos-Choline-8 ~5 minutes before grid preparation. Quantifoil R1.2/1.3 400 mesh holey carbon Au grids were glow discharged for 20s, and 3.5 *μL* of sample was applied, incubated for 5 minutes at 20°C and 100% humidity, and manually blotted from below. An additional 3.5 *μL* of sample was applied and after 30 seconds of incubation, grids were blotted for 3s with a blot force of 0, and plunge frozen in liquid ethane using a FEI Vitrobot Mark IV. For data acquisition, grids were loaded onto a 300-kV Titan Krios transmission electron microscope with a Gatan K3 Summit direct electron detector and a GIF quantum energy filter with a slit width of 20 eV and 5,448 movies were collected in superresolution mode with a pixel size of 0.435 Å and a defocus range of 1.5 to 2.5 μm using SerialEM (Mastronarde, 2005). The movies were recorded with 50 frames, a 1.5 second total exposure time (0.05s/frame) and a dose rate of 25 e^−^/pix/s which gave a cumulative dose of 50.7 e^−^/Å^2^ (1.01 e^−^/Å^2^/frame).

For *PLCβ*3 and reconstituted liposomes with *Gβγ*, *PLCβ*3 and liposomes with 1:15 (wt/wt) *Gβγ* were mixed at final concentrations of 0.5 *mg*/*mL* and 17.5 *mM* respectively and incubated at room temperature for one hour. The sample was supplemented with 3 *mM* of Fluorinated Fos-Choline-8 ~5 minutes before grid preparation. Quantifoil R1.2/1.3 400 mesh holey carbon Au grids were glow discharged for 20s, and 3.5 *μL* of sample was applied, incubated for 5 minutes at 22°C and 100% humidity, and manually blotted from below. An additional 3.5 *μL* of sample was applied and after 30 seconds of incubation, grids were blotted for 3.5s with a blot force of 0 and plunge frozen in liquid ethane using a FEI Vitrobot Mark IV. For data acquisition, grids were loaded onto a 300-kV Titan Krios transmission electron microscope, located at the HHMI Janelia Research Campus, with a Gatan K3 Summit direct electron detector and a GIF quantum energy filter with a slit width of 20 eV and 27,454 movies were collected in superresolution mode with a pixel size of 0.4195 Å and a defocus range of 1.5 to 2.5 *μm* using SerialEM (Mastronarde, 2005). The movies were recorded with 50 frames, a 4.194 second total exposure time (0.084s/frame) and a total dose of 60 e^−^/Å^2^ (1.2 e^−^/Å^2^/frame).

For *PLCβ*3 and *Gβγ* reconstituted into nanodiscs, nanodiscs were concentrated to 4 *mg*/*mL* using a 0.5-mL Amicon concentrator with 100-kDa molecular weight cutoff, mixed with *PLCβ*3 to final concentration of 3.5 *mg*/*mL* nanodiscs and 2.5 *mg*/*mL PLCβ* and incubated at room temperature for one hour. The sample was supplemented with 3 *mM* of Fluorinated Fos-Choline-8 ~5 minutes before grid preparation. Quantifoil R1.2/1.3 400 mesh holey carbon Au grids were glow discharged for 20s, and 3.5 *μL* of sample was applied. After 20s incubation at 16°C and 100% humidity, grids were blotted for 2.5s with a blot force of −3, and plunge frozen in liquid ethane using a FEI Vitrobot Mark IV. For data acquisition, grids were loaded onto a 300-kV Titan Krios transmission electron microscope with a Gatan K3 Summit direct electron detector and a GIF quantum energy filter with a slit width of 20 eV and 25,063 movies were collected in counting mode with a pixel size of 0.844 Å and a defocus range of 1 to 2.5 μm using Leginon (Suloway et al., 2005). The movies were recorded with 40 frames, a 2 second total exposure time (0.05s/frame) and total dose of 69.14 e^−^/Å^2^ (1.72 e^−^/Å^2^/frame).

#### Cryo-EM data processing

For *PLCβ*3 in solution without liposomes, motion correction was performed using MotionCorr 2 with 2x binning via Relion (in Relion 3.1) and contrast transfer function (CTF) estimation was carried out using CTFfind4 (Rohou and Grigorieff, 2015; Zheng et al., 2017; Zivanov et al., 2018). 1,489,175 particles were picked using Laplacian-of-Gaussian (LoG) picking in Relion 3.1, extracted without binning with a box size of 211.7 Å, and imported into cyroSPARC (Punjani et al., 2017; Zheng et al., 2017). 505,209 particles were selected after several rounds of 2D classification and used to generate an initial model (*ab initio* reconstruction), which resembled the crystal structure of PLCβ3 with density for both the catalytic core and the distal CTD. 195,277 particles were selected from heterogenous refinement using the PLCβ3 initial model and two junk models and imported into Relion 3.1using the csparc2star.py script (Asarnow et al., 2019) for an additional round of 3D classification with global alignment, which yielded a subset of 136,521 particles. These particles were subjected to 2D classification in Relion ignoring the CTF until the first peak to generate templates for auto-picking. Micrographs with an estimated resolution worse than 7 Å were discarded and the remaining 2,477 were used for template-based picking resulting in 1,021,849 particles, which were extracted with a box size of 211.7 Å. These particles were sorted with several rounds of 2D classification in cryoSPARC2 resulting in 771,044 particles, which were subjected to *ab initio* reconstruction with 5 classes: two junk classes, two classes resembling the PLCβ3 crystal structure with density for the catalytic core and the distal CTD, and one class resembling the PLCβ3 catalytic core. These particles were then subjected to heterogenous refinement with five models including one junk model, three models containing density for both the catalytic core and distal CTD and one containing only density for the catalytic core. 263,592 particles were sorted into the class with just the catalytic core, imported into Relion 3.1 using the csparc2star.py script (Asarnow et al., 2019), and refined to 4.3 Å. The remaining particles yielded low resolution reconstructions of PLCβ3 with varying amounts of density for the catalytic core and distal CTD. Extensive 3D classification with and without alignment including focused classification did not improve reconstructions with both domains, even with a larger box size of 276.5 Å. The 263,592 particles in the catalytic core reconstruction were subjected to two rounds of Bayesian particle polishing (Zivanov et al., 2018) and the resolution improved to 4 Å. To ensure that box size was not too small to accommodate the distal CTD, these particles were re-extracted with a box size of 276.5 Å and subjected to an additional three rounds of Bayesian particle polishing (Zivanov et al., 2019) and one round of CTF refinement (Zivanov et al., 2018) resulting in a 3.8 Å reconstruction. Attempts to recover density for the distal CTD from the reconstruction were unsuccessful. To improve the resolution, one round of masked 3D classification without alignment was carried out, resulting in the selection of 67,716 particles which refined to 3.6 Å. Resolution was evaluated using the 0.143 criterion of the Fourier shell correlation trough the PostProcess job in Relion (Scheres and Chen, 2012) (Figure S3). This map was used for model building.

For *PLCβ*3 alone on vesicles, motion correction was performed using MotionCorr 2 with 2x binning via Relion (in Relion 3.1) and contrast transfer function (CTF) estimation was carried out using CTFfind4 (Rohou and Grigorieff, 2015; Zheng et al., 2017; Zivanov et al., 2018). ~1,100 particles were manually picked from 23 micrographs in crYOLO (Wagner et al., 2019) and used to train a new model for picking (Figure S4A-C). 244,212 particles were picked, manually inspected to ensure integrity of picks with the new model, extracted with 2x binning and a box size of 222.7 Å in Relion 3.1, and imported into cryoSPARC (Punjani et al., 2017; Zivanov et al., 2018). 190,186 particles were selected after two rounds of sorting with 2D classification and subjected to *ab initio* reconstruction with three classes, one of which resembled *PLCβ*3 with density for both the distal CTD and the catalytic core. Particles were sorted with a resolution cutoff of 5 Å (206,044 particles remaining) and refined to this model to attempt to align the membranes yielding the reconstruction shown in Figure S4C. The particles were subjected to extensive 2D and 3D classification with and without alignment in Relion 3.1 and cryoSPARC, with and without signal subtraction of the membrane, and focused classification on the individual domains but no higher resolution reconstructions were obtained (Figure S4).

For the *PLCβ*3/*Gβγ* complex on liposomes, motion correction was performed with 2x binning using the Relion implementation (in Relion 3.1) and CTF estimation was carried out using CTFfind4 (Rohou and Grigorieff, 2015; Zheng et al., 2017; Zivanov et al., 2018). 2,411,476 particles were picked using the model trained for *PLCβ*3 on vesicles in crYOLO (Wagner et al., 2019) and sorted with an estimated resolution cutoff of 5.5 Å, resulting in 1,997,424 particles. A subset of 416,198 particles from 3,516 micrographs were extracted with 2x binning and a box size of 214.8 Å and imported into cryoSPARC3 for initial analysis (Punjani et al., 2017; Zivanov et al., 2018). 337,132 particles were selected from sorting with 2D classification and subjected to *ab initio* reconstruction with four classes. One class with 66,007 particles yielded a low resolution reconstruction showing density for the membrane and external protein. All 416,198 particles were refined to this model to attempt to align the membranes, imported into Relion 3.1 using the csparc2star.py script (Asarnow et al., 2019) and subjected to 2D classification without alignment ignoring the CTF until the first peak. 32,673 particles from 2D averages showing two features on the membrane surface were selected, imported into cryoSPARC3 and subjected to *ab initio* reconstruction with two classes (Figure S5). One class yielded a model with membrane density and two external features. All 1,997,424 particles were extracted with 2x binning and a box size of 255 Å and refined to this model in Relion to attempt to align the membranes. The membrane density was subtracted and the particles were subjected to several rounds of 3D classification. ~400,000 particles populated classes that resembled the *PLCβ*3 catalytic core bound to two *Gβγ*s and 121,394 particles from one class produced a reconstruction with clear secondary structure features. These particles were reverted to the original coordinates to undo the signal subtraction, re-extracted without binning and a box size of 289 Å, and imported into cryoSPARC3 for further analysis. Non-uniform refinement yielded a 4.4 Å reconstruction and included strong density for the membrane. Local refinement with a mask on the protein complex yielded a 3.9 Å reconstruction of the protein complex. These particles were subjected to Bayesian particle polishing and CTF refinement in Relion 3.1 (Zivanov et al., 2018; Zivanov et al., 2019) and refined again in cryoSPARC3 resulting in a 3.8 Å reconstruction from non-uniform refinement and a 3.5 Å reconstruction from local refinement with a mask on the protein complex. The resolution was evaluated using the 0.143 criterion of the Fourier shell correlation in cryoSPARC3 (Scheres and Chen, 2012). The map from local refinement was used for model building. These particles were subjected to additional 2D and 3D classification without alignment in Relion to investigate the position of the complex on the membrane. Resulting 3D classes were refined in cryoSPARC3 using non-uniform refinement and used for subsequent analysis (Figure 7, S5).

For the *PLCβ* ∙ *Gβγ* complex on nanodiscs, motion correction was performed using the Relion implementation (in Relion 3.1) and CTF estimation was carried out using CTFfind4 (Rohou and Grigorieff, 2015; Zheng et al., 2017; Zivanov et al., 2018). 3,993,932 particles were picked using crYOLO with the general model (Wagner et al., 2019), imported into Relion 3.1 and extracted 2x binned with a box size of 324 Å. Particles were sorted with a 5 Å estimated resolution cutoff in Relion resulting in 2,845,571 particles, which were imported into cryoSPARC3 and further sorted with 2D classification (Punjani et al., 2017; Zivanov et al., 2018). An initial model was determined using 1,166,729 particles following 2D classification, which resembled the *PLCβ*3 catalytic core bound to two *Gβγ*s. These particles were refined to the initial model resulting in a low resolution reconstruction. Focused 3D classification on the *PLCβ* catalytic core and *Gβγ* 2 was carried out in Relion 3.1, which yielded a reconstruction with 947,408 particles with secondary structure features. The model was improved by iterative 3D classification in Relion 3.1 and homogenous refinement in cryoSPARC3, which yielded a ~3.5 Å reconstruction from 55,623 particles. The map showed some artifacts from overfitting and over-sampled orientations. To improve the map quality and obtain more particles, sorting was repeated. The initial 2,845,571 particles were sorted to 2,428,922 particles with one round of 2D classification to remove obvious junk and subjected to heterogenous refinement in cryoSPARC3 with the map of the PLCβ3/Gβγ complex and two junk maps. 828,450 particles were sorted into the *PLCβ*3/*Gβγ* complex class and refined to 4.2 Å. Iterative 3D classification with and without alignment in Relion 3.1 and cryoSPARC4 yielded a subset of 168,057 particles which refined to 3.8 Å using non-uniform refinement with improved map quality. These particles were re-extracted without binning and underwent two rounds of Bayesian particle polishing in Relion (Zivanov et al., 2019) resulting in a 3.3 Å reconstruction from cryoSPARC3 non-uniform refinement. A final round of 3D classification without alignment in Relion resulted in a subset of 53,984 particles, which yielded a 3.3 Å reconstruction via local refinement in cryoSPARC3 with a mask on the protein complex. The resolution was evaluated using the 0.143 criterion of the Fourier shell correlation in cryoSPARC (Scheres and Chen, 2012). This map was free of artifacts and was used for model building.

#### Model building and Validation

For *PLCβ*3 in solution without liposomes, the crystal structure of *PLCβ*3 catalytic core from the PDB 4GNK (Lyon et al., 2013) was used as a starting model. It was fit into the density and refined with PHENIX real-space refine (Afonine et al., 2013) and manually inspected and adjusted where necessary. Regions with poor or weak density were removed. The proximal CTD (from 867 to 881) was excised from the crystal structure and fit into its density. The final model contains residues 12-92, 97-470, 574-850, and 867-881 and the side chains of E14, R169, K184, E191, K196, R199, E201, E232, K238, D777 were removed due to lack of density.

For the *PLCβ*3/*Gβγ* complex on liposomes, the model for *PLCβ*3 in solution was merged with two copies of *Gβγ* from PDB 4KFM (Whorton and MacKinnon, 2011) to use as the starting model. It was fit into the density and refined with PHENIX real-space refine (Afonine et al., 2013) and manually inspected and adjusted where necessary. Regions with poor or weak density were removed. The final model consists of *PLCβ*3: 13-92, 97-470, 576-849, 867-881 with the side chains of E14, E88, K184, E187, E191, R199, E201, R288, E303, R369, E373, E386, K603, R609, K631, K641, D777, K761, R872, and R874 removed due to lack of density, *Gβ* 1: 4-126, and 133-340 with the side chains of R137, R214, E215, and R256 removed due to lack density, *Gγ* 1: 8-62, *Gβ* 2: 2-126, and 133-340, with side chains of D292 and R52 removed due to lack of density, *Gγ* 2: 8-51.

For the *PLCβ*3/*Gβγ* complex on nanodiscs, the model for the complex on liposomes was used as the starting mode. It was fit into the density and refined with PHENIX real-space refine (Afonine et al., 2013) and manually inspected and adjusted where necessary. Regions with poor or weak density were removed. The final model consists of *PLCβ*3: 13-92, 97-470, 576-849, 867-881 with the sidechains of E14, D71, K82, E88, K184, E303, E373, D777 removed due to lack of density, *Gβ* 1: 4-126, and 133-340 with sidechains removed from R214, E215, M216, Q259 due to lack of density, *Gγ* 1: 8-62, *Gβ* 2: 2-126, and 133-140 with sidechains removed from : D153, Q175, R191, R256 due to poor density, and *Gγ* 2: 8-51. Model quality was assessed with validation in Phenix using MolProbity score (Chen et al., 2010) and geometry evaluation. Figures were made using ChimeraX (Pettersen et al., 2004; Pettersen et al., 2021).

For all chains, sidechains were not removed in areas of week density due to prior knowledge of structure of each component.

### Key Resources Table

| REAGENT or RESOURCE | SOURCE | IDENTIFIER |
| --- | --- | --- |
| Chemicals, peptides, and recombinant proteins |  |  |
| Sodium cholate hydrate | Sigma | Cat# C6445 |
| Bio-Beads SM-2 Resin | Bio-rad | Cat# 1523920 |
| n-Decyl-β-D-Maltopyranoside | anatrace | Cat# D322S |
| n-Dodecyl-β-D-Maltopyranoside | anatrace | Cat# D310S |
| Fos-Choline-8, Fluorinated | anatrace | Cat# F300F |
| LD655-maleimide | Lumidyne technologies | Cat# 09 |
| 1,2-dioctanoyl-sn-glycero-3-phospho-(1'-myo-inositol-4',5'-bisphosphate) (C8-PI(4,5)P<sub>2</sub>) | Avanti polar lipids | Cat# 850185 |
| 1-palmitoyl-2-oleoyl-sn-glycero-3-phosphoethanolamine (POPE) | Avanti polar lipids | Cat# 850757C |
| 1-palmitoyl-2-oleoyl-sn-glycero-3-phospho-(1'-rac-glycerol) (sodium salt) (POPG) | Avanti polar lipids | Cat# 840457C |
| 1,2-dioleoyl-sn-glycero-3-phosphoethanolamine (DOPE) | Avanti polar lipids | Cat# 850725C |
| 1-palmitoyl-2-oleoyl-glycero-3-phosphocholine (POPC) | Avanti polar lipids | Cat# 850457C |
| 1-palmitoyl-2-oleoyl-sn-glycero-3-phospho-L-serine (sodium salt) (POPS) | Avanti polar lipids | Cat# 840034C |
| 1,2-dioleoyl-sn-glycero-3-phosphoethanolamine-N-(lissamine rhodamine B sulfonyl) (ammonium salt) (18:1 Liss Rhod PE) | Avanti polar lipids | Cat# 810150C |
| L-α-phosphatidylinositol-4,5-bisphosphate (Brain, Porcine) (ammonium salt) (Brain PI(4,5)P<sub>2</sub>) | Avanti polar lipids | Cat# 840046X |
| Penicillin-Streptomycin (10,000 U/mL) | Thermo Fisher | Cat# 15140122 |
| Cellfectin II Reagent | Thermo Fisher | Cat# 10362100 |
| Fetal bovine serum | Thermo Fisher | Cat# 16000044 |
| Penicillin-Streptomycin (10,000 U/mL) | Thermo Fisher | Cat# 15140122 |
| L-Glutamine (200 mM) | Thermo Fisher | Cat# 25030081 |
| FreeStyle 293 expression medium | Thermo Fisher | Cat# 12338018 |
| Express Five SFM | Thermo Fisher | Cat# 10486025 |
| Sf-900 II SFM | Thermo Fisher | Cat# 10902096 |
| Critical Commercial Assays |  |  |
| CNBR-activated Sepharose 4B | Cytiva | Cat# 17043002 |
| Superose 6 Increase 10/300 GL | Cytiva | Cat# 29091569 |
| Superdex 200 Increase 10/300 GL | Cytiva | Cat# 28990944 |
| KLD enzyme mix | NEB | Cat# M0554S |
| NEBuilder® HiFi DNA Assembly Master Mix | NEB | Cat# E2621S |
| Deposited data |  |  |
| *PLCβ3* catalytic core without membranes atomic model | This study | PDB: 8EMV |
| *PLCβ3* catalytic core without membranes density map | This study | EMD: EMD-28266 |
| *PLCβ3* catalytic core in complex with *Gβγ* on vesicles atomic model | This study | PDB: 8EMW |
| *PLCβ3* catalytic core in complex with *Gβγ* on vesicles density map (excluding membrane) | This study | EMD: EMD-28267 |
| *PLCβ3* catalytic core in complex with *Gβγ* on nanodiscs atomic model | This study | PDB: 8EMX |
| *PLCβ3* catalytic core in complex with *Gβγ* on nanodiscs density map | This study | EMD: EMD-28268 |
| Experimental models: Cell lines |  |  |
| *Homo sapiens*: HEK293S GnT1- | ATCC | Cat# CRL-3022 |
| *Spodoptera frugiperda*: Sf9 | ATCC | Cat# CRL-1711 |
| *Trichoplusia ni*: High Five | Thermo Fisher | Cat# B85502 |
| Recombinant DNA |  |  |
| h*PLCβ3* | Dr. John Sondek from Charpentier et al., 2014 | N/A |
| Gβ1 | (Whorton and MacKinnon, 2013) | N/A |
| Gγ2 | (Whorton and MacKinnon, 2013) | N/A |
| pEG BacMam | (Goehring et al., 2014) | N/A |
| Software and algorithms |  |  |
| SerialEM | (Mastronarde, 2005) | http://bio3d.colorado.edu/SerialEM |
| Leginon | (Suloway et al., 2005) | https://emg.nysbc.org/redmine/projects/leginon/wiki/Leginon_Home_page |
| MotionCor2 | (Zheng et al., 2017) | https://msg.ucsf.edu/software |
| CTFFind4 | (Rohou and Grigorieff, 2015) | https://grigoriefflab.umassmed.edu/ctffind4 |
| crYOLO | (Wagner et al., 2019) | https://cryolo.readthedocs.io/en/stable/ |
| cryoSPARC | (Punjani et al., 2017) | https://cryosparc.com/ |
| RELION-3.1 | (Zivanov et al., 2018) | https://github.com/3dem/relion |
| SBGrid | (Morin et al., 2013) Morin et al., 2018 | https://sbgrid.org/ |
| COOT | (Emsley et al., 2010) | https://www2.mrc-lmb.cam.ac.uk/personal/pemsley/coot/ |
| UCSF Chimera | (Pettersen et al., 2004) | https://www.cgl.ucsf.edu/chimera/download.html |
| UCSF ChimeraX | (Pettersen et al., 2021) | https://www.cgl.ucsf.edu/chimerax/download.html |
| PHENIX | (Afonine et al., 2013) | https://phenix-online.org/download/ |
| pClamp | N/A | https://support.moleculardevices.com/s/article/Axon-pCLAMP-10-Electrophysiology-Data-Acquisition-Analysis-Software-Download-Page |
| QtiPlot | N/A | https://www.qtiplot.com/download.html |
| Other |  |  |
| R1.2/1.3 400 mesh Au holey carbon grids | Quantifoil | 1210627 |

## References

1. Afonine, P.V., Headd, J.J., Terwilliger, T.C., and Adams, P.D. (2013). PHENIX News. Computational Crystallography Newsletter 4, 43–44.

2. Asarnow, D., Palovcak, E., and Cheng, Y. (2019). asarnow/pyem: UCSF pyem v0.5 (Zenodo).

3. Berridge, M.J. (1987). Inositol Trisphosphate and Diacylglycerol: Two Interacting Second Messengers. Annual Review of Biochemistry 56, 159–193.

4. Camps, M., Carozzi, A., Schnabel, P., Scheer, A., Parker, P.J., and Gierschik, P. (1992). Isozyme-selective stimulation of phospholipase C-β2 by G protein βγ-subunits. In Nature, pp. 684–686.

5. Carroll, R.C., Morielli, A.D., and Peralta, E.G. (1995). Coincidence detection at the level of phospholipase C activation mediated by the m4 muscarinic acetylcholine receptor. In Current Biology, pp. 536–544.

6. Charpentier, T.H., Waldo, G.L., Barrett, M.O., Huang, W., Zhang, Q., Harden, T.K., and Sondek, J. (2014). Membrane-induced allosteric control of phospholipase C-β isozymes. In Journal of Biological Chemistry, pp. 29545–29557.

7. Chen, V.B., Arendall, W.B., Headd, J.J., Keedy, D.A., Immormino, R.M., Kapral, G.J., Murray, L.W., Richardson, J.S., and Richardson, D.C. (2010). MolProbity: All-atom structure validation for macromolecular crystallography. Acta Crystallographica Section D: Biological Crystallography 66, 12–21.

8. Drin, G., Douguet, D., and Scarlata, S. (2006). The Pleckstrin homology domain of phospholipase Cβ transmits enzymatic activation through modulation of the membrane-domain orientation. In Biochemistry, pp. 5712–5724.

9. Emsley, P., Lohkamp, B., Scott, W.G., and Cowtan, K. (2010). Features and development of Coot. Acta Crystallographica Section D: Biological Crystallography 66, 486–501.

10. Fisher, I.J., Jenkins, M.L., Tall, G.G., Burke, J.E., and Smrcka, A.V. (2020). Activation of Phospholipase C β by Gβγ and Gαq Involves C-Terminal Rearrangement to Release Autoinhibition. In Structure, pp. 1–10.

11. Ford, C.E., Skiba, N.P., Bae, H., Daaka, Y., Reuveny, E., Shekter, L.R., Rosal, R., Weng, G., Yang, C.S., Iyengar, R., et al. (1998). Molecular basis for interactions of G protein βΓ subunits with effectors. Science 280, 1271–1274.

12. Goehring, A., Lee, C.H., Wang, K.H., Michel, J.C., Claxton, D.P., Baconguis, I., Althoff, T., Fischer, S., Garcia, K.C., and Gouaux, E. (2014). Screening and large-scale expression of membrane proteins in mammalian cells for structural studies. Nature Protocols 9, 2574–2585.

13. Goličnik, M. (2012). On the Lambert W function and its utility in biochemical kinetics. Biochemical Engineering Journal 63, 116–123.

14. Götzke, H., Kilisch, M., Martínez-Carranza, M., Sograte-Idrissi, S., Rajavel, A., Schlichthaerle, T., Engels, N., Jungmann, R., Stenmark, P., Opazo, F., et al. (2019). The ALFA-tag is a highly versatile tool for nanobody-based bioscience applications. Nature Communications 10, 1–12.

15. Grinkova, Y.V., Denisov, I.G., and Sligar, S.G. (2010). Engineering extended membrane scaffold proteins for self-assembly of soluble nanoscale lipid bilayers. Protein Engineering, Design and Selection 23, 843–848.

16. Gutman, O., Walliser, C., Piechulek, T., Gierschik, P., and Henis, Y.I. (2010). Differential regulation of phospholipase C-β2 activity and membrane interaction by Gαq, Gβ1γ 2, and Rac2. In Journal of Biological Chemistry, pp. 3905–3915.

17. Hansen, S.B. (2015). Lipid agonism: The PIP2 paradigm of ligand-gated ion channels. In Biochimica et Biophysica Acta - Molecular and Cell Biology of Lipids (Elsevier B.V.), pp. 620–628.

18. Hicks, S.N., Jezyk, M.R., Gershburg, S., Seifert, J.P., Harden, T.K., and Sondek, J. (2008). General and Versatile Autoinhibition of PLC Isozymes. In Molecular Cell, pp. 383–394.

19. Illenberger, D., Schwald, F., Pimmer, D., Binder, W., Maier, G., Dietrich, A., and Gierschik, P. (1998). Stimulation of phospholipase C-β2 by the Rho GTPases Cdc42Hs and Rac1. The EMBO Journal 17, 6241–6249.

20. Illenberger, D., Walliser, C., Strobel, J., Gutman, O., Niv, H., Gaidzik, V., Kloog, Y., Gierschik, P., and Henis, Y.I. (2003). Rac2 regulation of phospholipase C-β2 activity and mode of membrane interactions in intact cells. In Journal of Biological Chemistry, pp. 8645–8652.

21. Jenco, J.M., Becker, K.P., and Morris, A.J. (1997). Membrane-binding properties of phospholipase C-β1 and phospholipase C-β: Role of the C-terminus and effects of polyphosphoinositides, G-proteins and Ca2+. In Biochemical Journal, pp. 431–437.

22. Jezyk, M.R., Snyder, J.T., Gershberg, S., Worthylake, D.K., Harden, T.K., and Sondek, J. (2006). Crystal structure of Rac1 bound to its effector phospholipase C-β2. Nature Structural & Molecular Biology 13, 1135–1140.

23. Kadamur, G., and Ross, E.M. (2013). Mammalian Phospholipase C. In Annual Review of Physiology, pp. 127–154.

24. Kadamur, G., and Ross, E.M. (2016). Intrinsic Pleckstrin Homology (PH) domain motion in phospholipase C-β exposes a Gβγ protein binding site. In Journal of Biological Chemistry, pp. 11394–11406.

25. Katz, A., Wu, D., and Simon, M.I. (1992). Subunits βγ of heterotrimeric G protein activate β2 isoform of phospholipase C. In Nature, pp. 686–689.

26. Lyon, A.M., Begley, J.A., Manett, T.D., and Tesmer, J.J.G. (2014a). Molecular mechanisms of phospholipase C β3 autoinhibition. In Structure, pp. 1844–1854.

27. Lyon, A.M., Dutta, S., Boguth, C.A., Skiniotis, G., and Tesmer, J.J.G. (2013). Full-length Gαq-phospholipase C-β3 structure reveals interfaces of the C-terminal coiled-coil domain. In Nature Structural and Molecular Biology (Nature Publishing Group), pp. 355–362.

28. Lyon, A.M., Taylor, V.G., and Tesmer, J.J.G. (2014b). Strike a pose: Gαq complexes at the membrane. In Trends in Pharmacological Sciences (Elsevier Ltd), pp. 23–30.

29. Lyon, A.M., and Tesmer, J.J.G. (2013). Structural insights into phospholipase C-β function. In Molecular Pharmacology, pp. 488–500.

30. Lyon, A.M., Tesmer, V.M., Dhamsania, V.D., Thal, D.M., Gutierrez, J., Chowdhury, S., Suddala, K.C., Northup, J.K., and Tesmer, J.J.G. (2011). An autoinhibitory helix in the C-terminal region of phospholipase C-β mediates Gαq activation. In Nature Structural and Molecular Biology (Nature Publishing Group), pp. 999–1005.

31. Mandal, K. (2020). Review of PIP2 in Cellular Signaling, Functions and Diseases. IJMS 21, 8342.

32. Mastronarde, D.N. (2005). Automated electron microscope tomography using robust prediction of specimen movements. Journal of Structural Biology 152, 36–51.

33. McLaughlin, S., Wang, J., Gambhir, A., and Murray, D. (2002). PIP2 and Proteins: Interactions, Organization, and Information Flow. Annual Review of Biophysics and Biomolecular Structure 31, 151–175.

34. Miller, C. (1983). Integral Membrane Channels: Studies in Model Membranes. Physiological Reviews 63, 1209–1242.

35. Morin, A., Eisenbraun, B., Key, J., Sanschagrin, P.C., Timony, M.A., Ottaviano, M., and Sliz, P. (2013). Collaboration gets the most out of software. eLife 2, e01456.

36. Nichols, C.G., and Lee, S.-j. (2018). Polyamines and potassium channels: A 25-year romance. Journal of Biological Chemistry 293, 18779–18788.

37. Okajima, F., Sato, K., Sho, K., and Kondo, Y. (1989). Stimulation of adenosine receptor enhances α1 -adrenergic receptor-mediated activation of phospholipase C and Ca2+ mobilization in a pertussis toxin-sensitive manner in FRTL-5 thyroid cells. In FEBS Letters, pp. 145–149.

38. Pettersen, E.F., Goddard, T.D., Huang, C.C., Couch, G.S., Greenblatt, D.M., Meng, E.C., and Ferrin, T.E. (2004). UCSF Chimera - A visualization system for exploratory research and analysis. Journal of Computational Chemistry 25, 1605–1612.

39. Pettersen, E.F., Goddard, T.D., Huang, C.C., Meng, E.C., Couch, G.S., Croll, T.I., Morris, J.H., and Ferrin, T.E. (2021). UCSF ChimeraX: Structure visualization for researchers, educators, and developers. Protein Science 30, 70–82.

40. Philip, F., Kadamur, G., Silos, R.G., Woodson, J., and Ross, E.M. (2010). Synergistic activation of phospholipase C-β3 by Gαq and Gβγ describes a simple two-state coincidence detector. In Current Biology, pp. 1327–1335.

41. Poveda, J.A., Giudici, A.M., Renart, M.L., Molina, M.L., Montoya, E., Fernández-Carvajal, A., Fernández-Ballester, G., Encinar, J.A., and González-Ros, J.M. (2014). Lipid modulation of ion channels through specific binding sites. Biochimica et Biophysica Acta - Biomembranes 1838, 1560–1567.

42. Punjani, A., Rubinstein, J.L., Fleet, D.J., and Brubaker, M.A. (2017). CryoSPARC: Algorithms for rapid unsupervised cryo-EM structure determination. Nature Methods 14, 290–296.

43. Rebres, R.A., Roach, T.I.A., Fraser, I.D.C., Philip, F., Moon, C., Lin, K.M., Liu, J., Santat, L., Cheadle, L., Ross, E.M., et al. (2011). Synergistic Ca2+ responses by Gαi- and Gαq-coupled G-protein-coupled receptors require a single PLCβ isoform that is sensitive to both Gβγ and Gαq. In Journal of Biological Chemistry, pp. 942–951.

44. Roach, T.I.A., Rebres, R.A., Fraser, I.D.C., DeCamp, D.L., Lin, K.M., Sternweis, P.C., Simon, M.I., and Seaman, W.E. (2008). Signaling and cross-talk by C5a and UDP in macrophages selectively use PLCβ3 to regulate intracellular free calcium. In Journal of Biological Chemistry, pp. 17351–17361.

45. Robinson, C.V., Rohacs, T., and Hansen, S.B. (2019). Tools for Understanding Nanoscale Lipid Regulation of Ion Channels. Trends in Biochemical Sciences 44, 795–806.

46. Rohou, A., and Grigorieff, N. (2015). CTFFIND4: Fast and accurate defocus estimation from electron micrographs. Journal of Structural Biology 192, 216–221.

47. Romoser, V., Ball, R., and Smrcka, A.V. (1996). Phospholipase C β2 Association with Phospholipid Interfaces Assessed by Fluorescence Resonance Energy Transfer. In Journal of Biological Chemistry, pp. 25071–25078.

48. Runnels, L.W., Jenco, J., Morris, A., and Scarlata, S. (1996). Membrane binding of phospholipases C-β1 and C-β2 is independent of phosphatidylinositol 4,5-bisphosphate and the αand βγ subunits of G proteins. In Biochemistry, pp. 16824–16832.

49. Scheres, S.H.W., and Chen, S. (2012). Prevention of overfitting in cryo-EM structure determination. Nature Methods 9, 853–854.

50. Shah, B.H., Siddiqui, A., Qureshi, K.A., Khan, M., Rafi, S., Ujan, V.A., Yaqub, Y., Rasheed, H., and Saeed, S.A. (2009). Co-activation of Gi and Gq proteins exerts synergistic effect on human platelet aggregation through activation of phospholipase C and Ca 2+ signalling pathways (Experimental and Molecular Medicine (1999) 31 (42-46)). In Experimental and Molecular Medicine, pp. 686.

51. Smrcka, A.V., and Sternweis, P.C. (1993). Regulation of purified subtypes of phosphatidylinositol-specific phospholipase C β by G protein α and βγ subunits. In Journal of Biological Chemistry, pp. 9667–9674.

52. Suloway, C., Pulokas, J., Fellmann, D., Cheng, A., Guerra, F., Quispe, J., Stagg, S., Potter, C.S., and Carragher, B. (2005). Automated molecular microscopy: The new Leginon system. Journal of Structural Biology 151, 41–60.

53. Wagner, T., Merino, F., Stabrin, M., Moriya, T., Antoni, C., Apelbaum, A., Hagel, P., Sitsel, O., Raisch, T., Prumbaum, D., et al. (2019). SPHIRE-crYOLO is a fast and accurate fully automated particle picker for cryo-EM. Communications Biology 2, 1–13.

54. Waldo, G.L., Ricks, T.K., Hicks, S.N., Cheever, M.L., Kawano, T., Tsuboi, K., Wang, X., Montell, C., Kozasa, T., Sondek, J., et al. (2010). Kinetic Scaffolding Mediated by a Phospholipase C–b and Gq Signaling Complex. In Science.

55. Wang, W., Touhara, K.K., Weir, K., Bean, B.P., and MacKinnon, R. (2016). Cooperative regulation by G proteins and Na+ of neuronal GIRK2 K+ channels. eLife 5, 1–15.

56. Wang, W., Whorton, M.R., and MacKinnon, R. (2014). Quantitative analysis of mammalian GIRK2 channel regulation by G proteins, the signaling lipid PIP2 and Na+ in a reconstituted system. In eLife, pp. e03671.

57. Ware, J.A., Smith, M., and Salzman, E.W. (1987). Synergism of platelet-aggregating agents. Role of elevation of cytoplasmic calcium. In Journal of Clinical Investigation, pp. 267–271.

58. White, S.H., Wimley, W.C., Ladokhin, A.S., and Hristova, K. (1998). [4] Protein folding in membranes: Determining energetics of peptide-bilayer interactions. In Methods in Enzymology (Academic Press), pp. 62–87.

59. Whorton, M.R., and MacKinnon, R. (2011). Crystal structure of the mammalian GIRK2 K + channel and gating regulation by G proteins, PIP 2, and sodium. Cell 147, 199–208.

60. Whorton, M.R., and MacKinnon, R. (2013). X-ray structure of the mammalian GIRK2-βγ G-protein complex. Nature 498, 190–197.

61. Zhao, C., and MacKinnon, R. (2022). Structural and functional analyses of a GPCR-inhibited ion channel TRPM3. Neuron *0*.

62. Zheng, Q., Jockusch, S., Zhou, Z., Altman, R.B., Zhao, H., Asher, W., Holsey, M., Mathiasen, S., Geggier, P., Javitch, J.A., et al. (2016). Electronic tuning of self-healing fluorophores for live-cell and single-molecule imaging. Chem Sci 8, 755–762.

63. Zheng, S.Q., Palovcak, E., Armache, J.P., Verba, K.A., Cheng, Y., and Agard, D.A. (2017). MotionCor2: Anisotropic correction of beam-induced motion for improved cryo-electron microscopy. Nature Methods 14, 331–332.

64. Zivanov, J., Nakane, T., Forsbeg, B., Kimanius, D., Hagen, W.J.H., Lindhal, E., and Scheres, S.H.W. (2018). RELION-3 : new tools for automated high-resolution cryo-EM structure determination. bioRxiv, 1–38.

65. Zivanov, J., Nakane, T., and Scheres, S.H.W. (2019). A Bayesian approach to beam-induced motion correction in cryo-EM single-particle analysis. IUCrJ 6, 5–17.

