## Supplementary material for "*Gβγ* Activates *PIP*2 Hydrolysis by Recruiting and Orienting *PLCβ* on the Membrane Surface": primary SI

| Collection Parameters | $PLC\beta3$ solution | $PLC\beta3/G\beta\gamma$ complex-liposomes | $PLC\beta3/G\beta\gamma$ complex-nanodiscs | $PLC\beta3$ liposomes |
|---|---|---|---|---|
| Accelerating Voltage (kV) | 300 | 300 | 300 | 300 |
| Number of frames | 40 | 50 | 40 | 50 |
| Dose ($e^-/Å^2$) | 42.87 | 60 | 69.14 | 50.7 |
| Defocus Range (μm) | 1-2.5 | 1.5-2.5 | 1-2.5 | 1.5-2.5 |
| Exposure Time (s) | 2 | 4.194 | 2 | 1.5 |
| Original Pixel size (Å) | 0.54 | 0.4195 | 0.844 | 0.435 |
| Map Parameters | | | | |
| Final Pixel Size (Å) | 1.08 | 0.839 | 0.844 | 0.87 |
| Symmetry | C1 | C1 | C1 | C1 |
| Total micrographs | 3,527 | 27,454 | 25,063 | 5,448 |
| Initial particles | 1,021,849 | 1,997,424 | 1,728,771 | 244,212 |
| Final particles | 67,716 | 121,394 | 53,984 | 244,212 |
| Map resolution (Å) | 3.59 | 3.47 | 3.33 | |
| FSC threshold | 0.143 | 0.143 | 0.143 | |
| Map local resolution range (Å) | 2.8-5.0 | 2.7-5.0 | 2.6-5.0 | |
| Map sharpening B factor ($Å^2$) | -111.735 | -116 | -79.1 | |
| Initial Model Used | 4GNK | 8EMV | 8EMV | |
| Model resolution (Å) ($FSC_{model}$=0.5) | 3.8 | 4.0 | 3.8 | |
| Model Composition | | | | |
| Nonhydrogen atoms | 5,949 | 11,732 | 11,735 | |
| Protein residues | 747 | 1506 | 1506 | |
| Ligands | 1 | 1 | 1 | |
| r.m.s. deviations bond length (Å) | 0.007 | 0.008 | 0.006 | |
| r.m.s. deviations bond length (Å) | 0.8 | 0.899 | 0.798 | |
| Validation | | | | |
| MolProbity Score | 1.53 | 1.77 | 1.59 | |
| Clash Score | 3.11 | 6.63 | 4.73 | |
| Poor Rotamers (%) | 0 | 0 | 0 | |
| Ramachandran Plot | | | | |
| Favored (%) | 93.50 | 93.88 | 94.95 | |
| Allowed (%) | 6.5 | 6.12 | 5.05 | |
| Disallowed (%) | 0 | 0 | 0 | |

Table S1: Cryo-EM collection parameters and model statistics. Related to Figure 4, 5, S3, S4, S5 and S7.

| Interface | Protein | Residues |
|---|---|---|
| $G\beta\gamma 1$ | $PLC\beta 3$ | R24, K27, I29, R38, N39, L40, P57, N58, M59, V89, R204, V123, Q166, D167, G168, and R169 |
| | $G\beta 1$ | K57, S98, W99, M101, L117, Y145, C204, D228, D267, N267, I270, C271, D290, D291, F292, N313, R314, and W332 |
| $G\beta\gamma 2$ | $PLC\beta 3$ | R185, T188, S192, R215, N218, K219, L222, P224, D227, L231, K236, G237, K238, P239, Y240, N282, Q284, F285, R288, M293, and E294 |
| | $G\beta 2$ | L55, A56, K57, Q75, D76, K78, S98, W99, M101, L117, T143, Y145, D186, M188, C204, D228, N230, D246, D290, N313, R314, W332, and D333 |

Table S2: Interface residues in the $PLC\beta 3/G\beta\gamma$ complex. Residues identified have buried surface area > 15 Å$^2$. W99, M101, L117, T143, D186, D228, W332 from $G\beta$ were shown to be important for $PLC\beta$ activation [1]. Related to Figure 5.

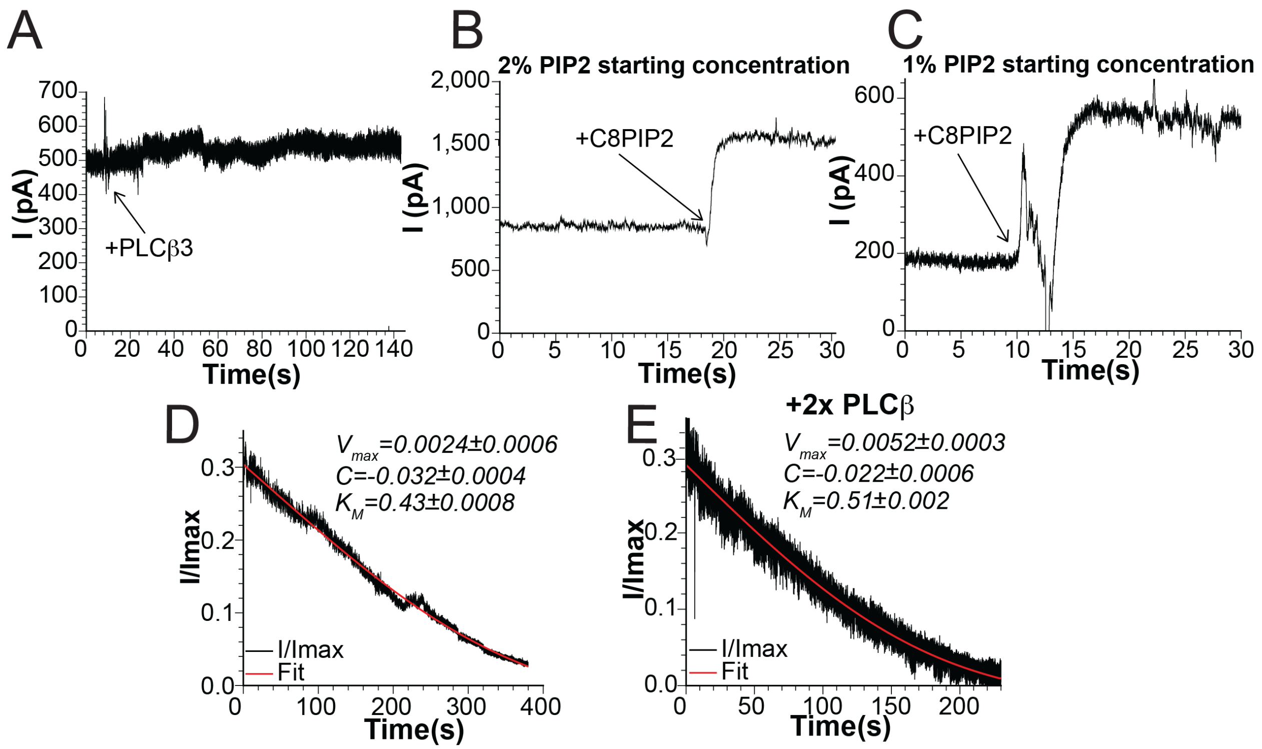


Figure S1 (Related to Figure 1-3): Planar lipid bilayer assay for $PLC\beta$ function. A: Representative current trace in the absence of $CaCl_2$ (2 mM EGTA) showing that addition of $PLC\beta3$ does not lead to current decay without $Ca^{2+}$, which is required for catalytic activity. B-C: Representative titration experiments for GIRK-ALFA with $PIP2$ starting with 2 mol% (B) or 1 mol% (C). C-D: normalized current decay in the absence of $G\beta\gamma$ with 1x (D) or 2x (E) $PLC\beta3$ concentration fit to equation 4 showing that $V_{max}$ depends on the added enzyme concentration. $R^2$=0.992 for D. $R^2$=0.969 E.

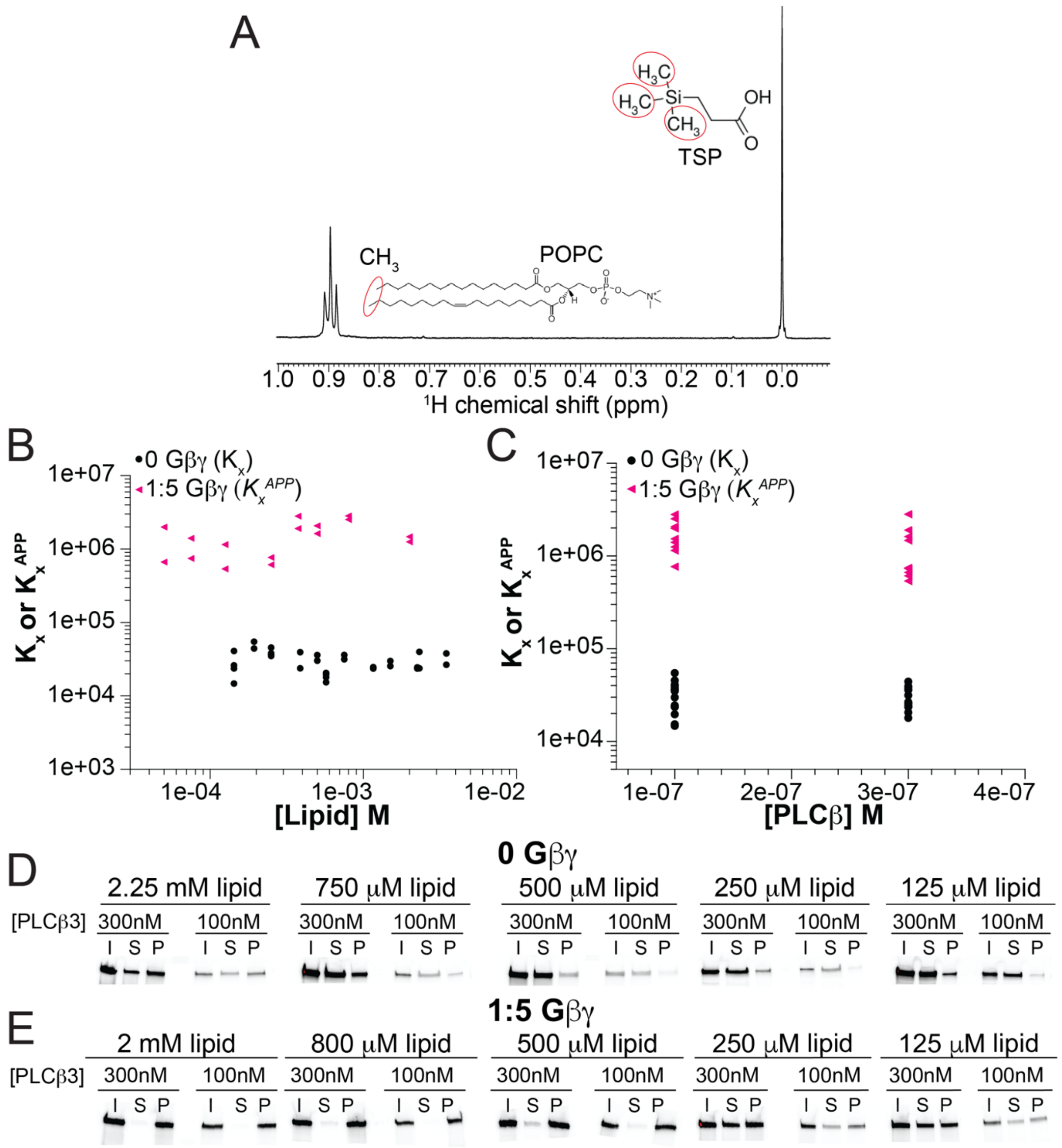


Figure S2 (Related to Figure 3): $G\beta\gamma$ enhanced membrane partitioning of $PLC\beta3$. A: representative $^{1}H$ NMR spectrum showing the lipid methyl peak at 0.9 ppm and the TSP peak at 0 ppm. Corresponding methyl groups are circled in red on the chemical structures. POPC is shown as a representative lipid. B-C: Plot of individual partition coefficients in the absence of Gβγ ($K_x$) or apparent partition coefficients in the presence of $G\beta\gamma$ ($K_x^{APP}$) determined for each experiment plotted against lipid concentration (B) or against $PLC\beta3$ concentration (C) showing that values do not vary with concentration of lipid (B) or $PLC\beta3$ (C). Experiments without $G\beta\gamma$ are shown as black spheres and experiments with $G\beta\gamma$ are shown as pink triangles. D-E: Example SDS-page gels imaged for LD655 fluorescence from binding experiments with (D) or without $G\beta\gamma$ (E). I represents input, S represents supernatant, and P represents pellet.

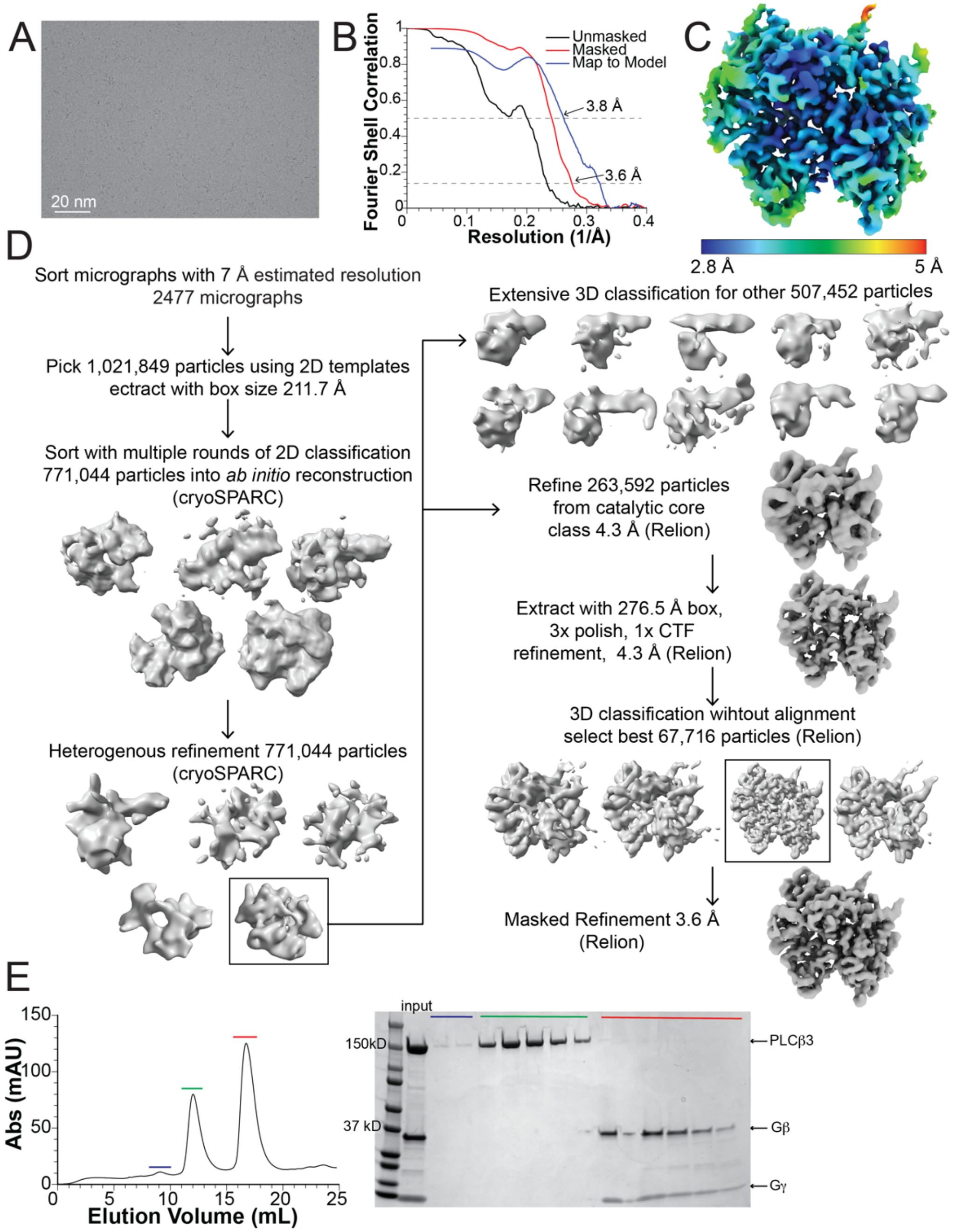


Figure S3 (related to Figure 4): Structure determination of $PLC\beta3$ in solution without membranes. A: representative micrograph. B: Fourier shell correlation (FSC) curves for the unmasked (black) and masked (red) maps and between the map and model (blue). The 0.143 and 0.5 thresholds are denoted by dashed lines. C: Final masked, sharpened map colored by local resolution determined by cryoSPARC. D: Summary of data processing steps, see methods. Maps shown are unsharpened. E: Attempt to form complex between $PLC\beta3$ and soluble $G\beta\gamma$ C68S. Proteins were mixed at a 1:2 ratio ($PLC\beta3$:$G\beta\gamma$) and ran on gel filtration using a Superdex 200 increase column (left). Peaks are color coded to SDS PAGE gel (right) showing $PLC\beta3$ and $G\beta\gamma$ do not comigrate.

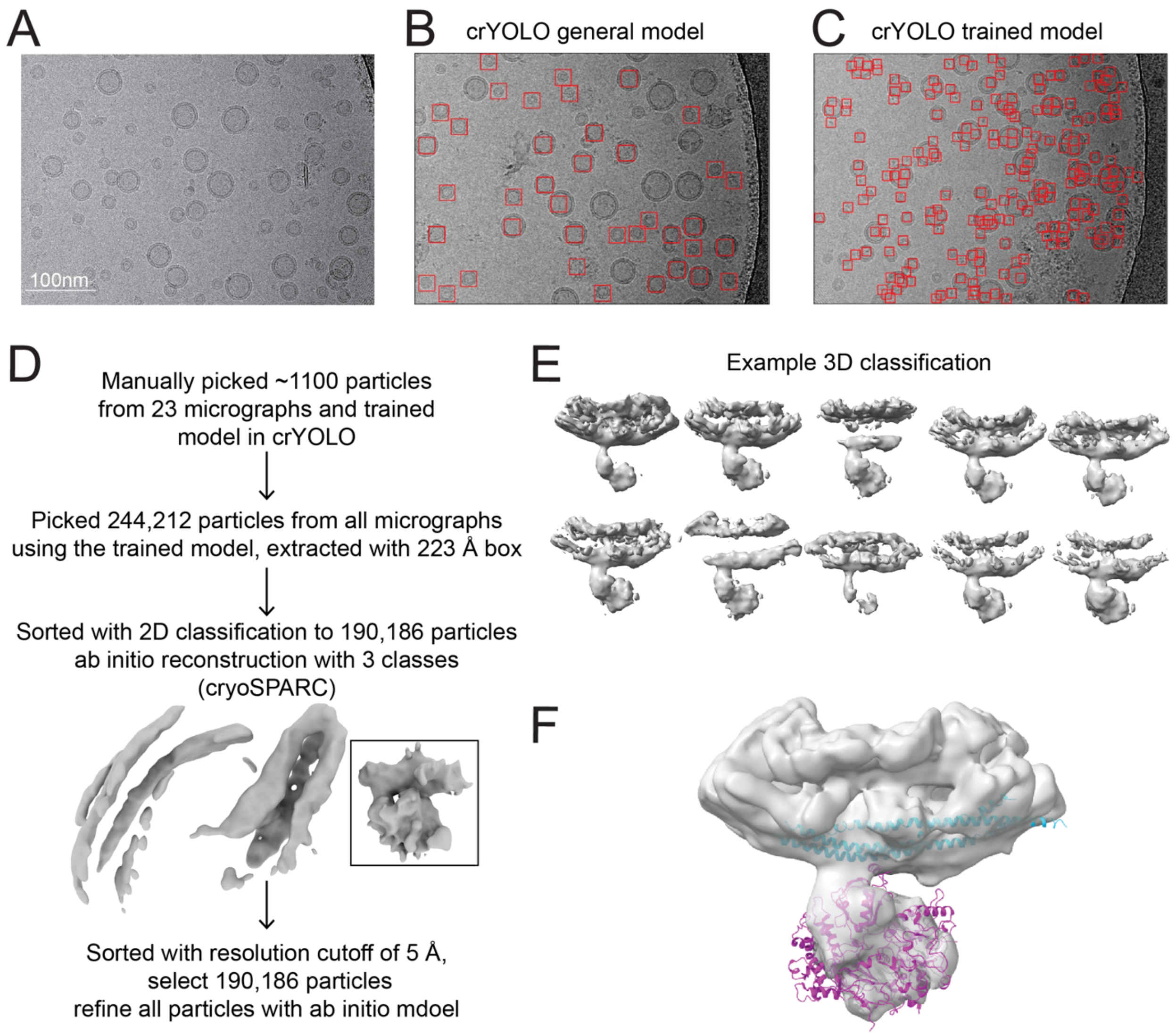


Figure S4 (related to Figure 4): Structure studies of $PLC\beta3$ associated with liposomes. A: representative micrograph. B-C: Example of particle picking using the crYOLO general model (C) or a crYOLO model trained on these micrographs (D) [2]. Picked particles are shown as a red box. D: Summary of data processing steps, see methods. E: Example of 3D classification of the final model run in cryoSPARC without alignment. F: Final unsharpened map with models for the distal CTD (blue, PDBID 4GNK [3]) and the catalytic core (pink) fit into the density. Extra density at the top is the membrane.

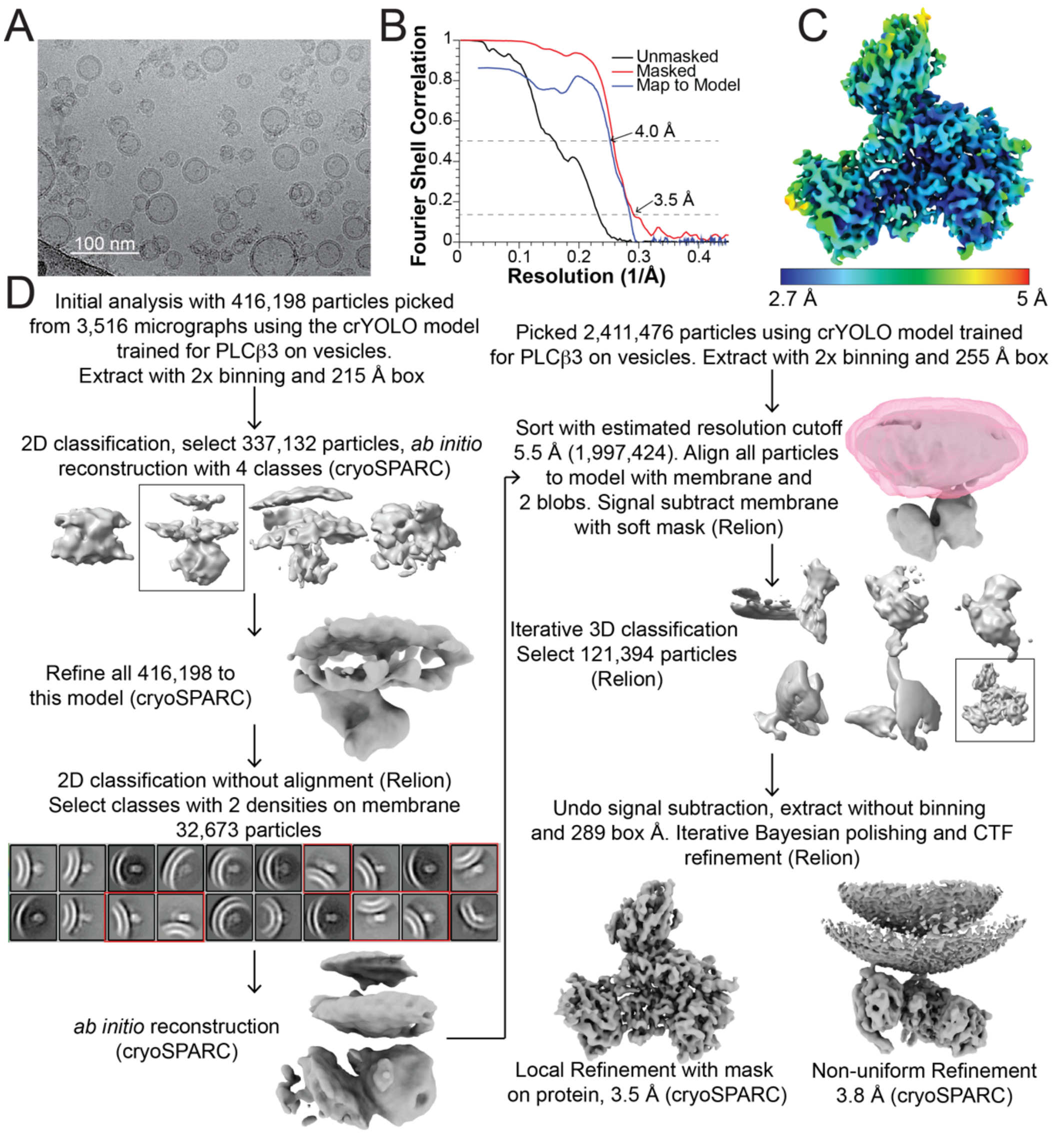


Figure S5 (related to Figure 5, 6): Structure determination of the $PLC\beta3 \cdot G\beta\gamma$ complex on liposomes comprised of 2DOPE:1POPC:1POPS. A: representative micrograph. B: Fourier shell correlation (FSC) curves for the unmasked (black) and masked (red) maps and between the map and model (blue). The 0.143 and 0.5 thresholds are denoted by dashed lines. C: Final masked, sharpened map colored by local resolution determined by cryoSPARC. D: Summary of data processing steps, see methods. Maps shown are unsharpened.

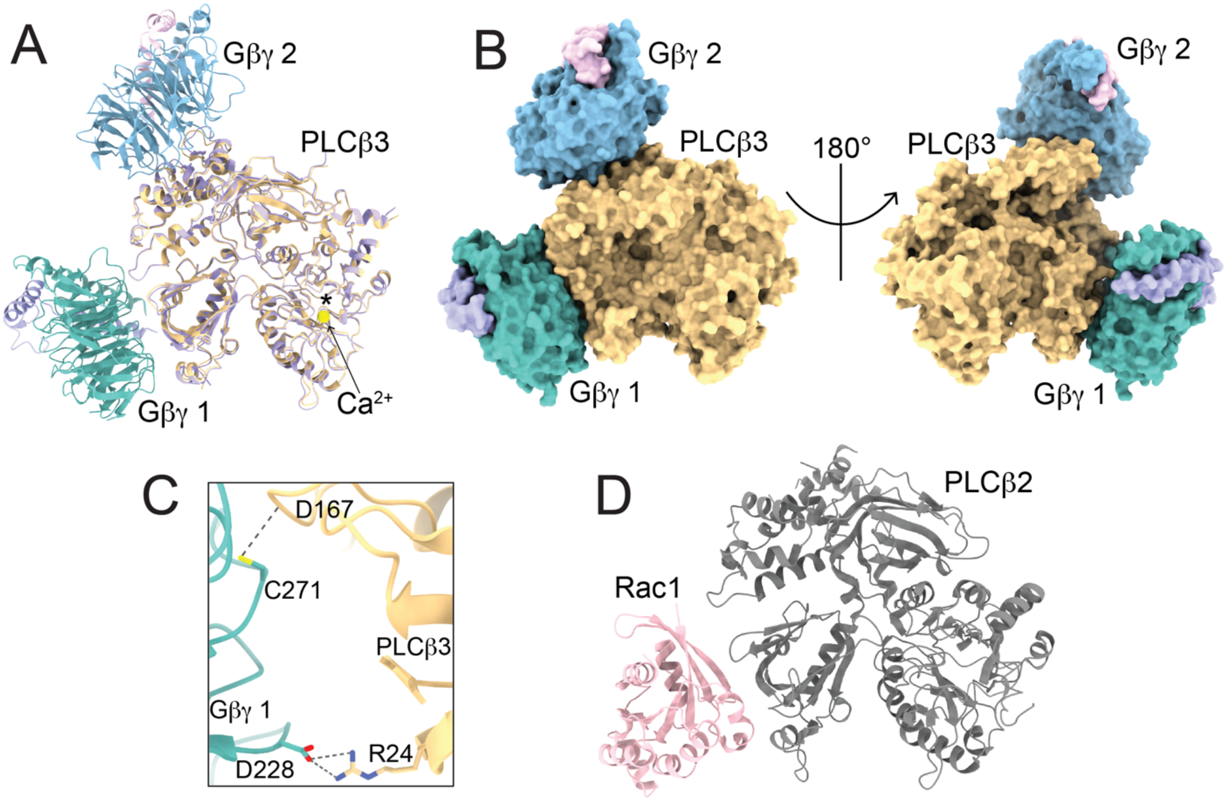


Figure S6 (related to Figure 5, 6): Interfaces of the $PLC\beta3 \cdot G\beta\gamma$ complex. A: Structural alignment of the $PLC\beta3$ catalytic core determined by cryo-EM (purple) and the $PLC\beta3 \cdot G\beta\gamma$ complex (colored by protein). RMSD ~0.7 Å. $PLC\beta3$ is yellow, $G\beta1$ is dark teal, $G\gamma1$ is light purple, $G\beta2$ is light blue and $G\gamma2$ is light pink. Calcium ion is shown as a yellow sphere and active site is denoted by asterisk. B: Surface representation of the $PLC\beta3 \cdot G\beta\gamma$ complex viewed from the top (left) and bottom (right) highlighting the $PLC\beta3$-$G\beta\gamma$ interfaces, colored as in A. C: Hydrogen bonds in the $G\beta\gamma1$ interface, $PLC\beta3$ is yellow and $G\beta$ is dark teal. D: $PLC\beta2$ catalytic core (gray) in complex with Rac1 (pink) highlighting similar positioning of $G\beta\gamma1$. PDBID-2FJU [4].

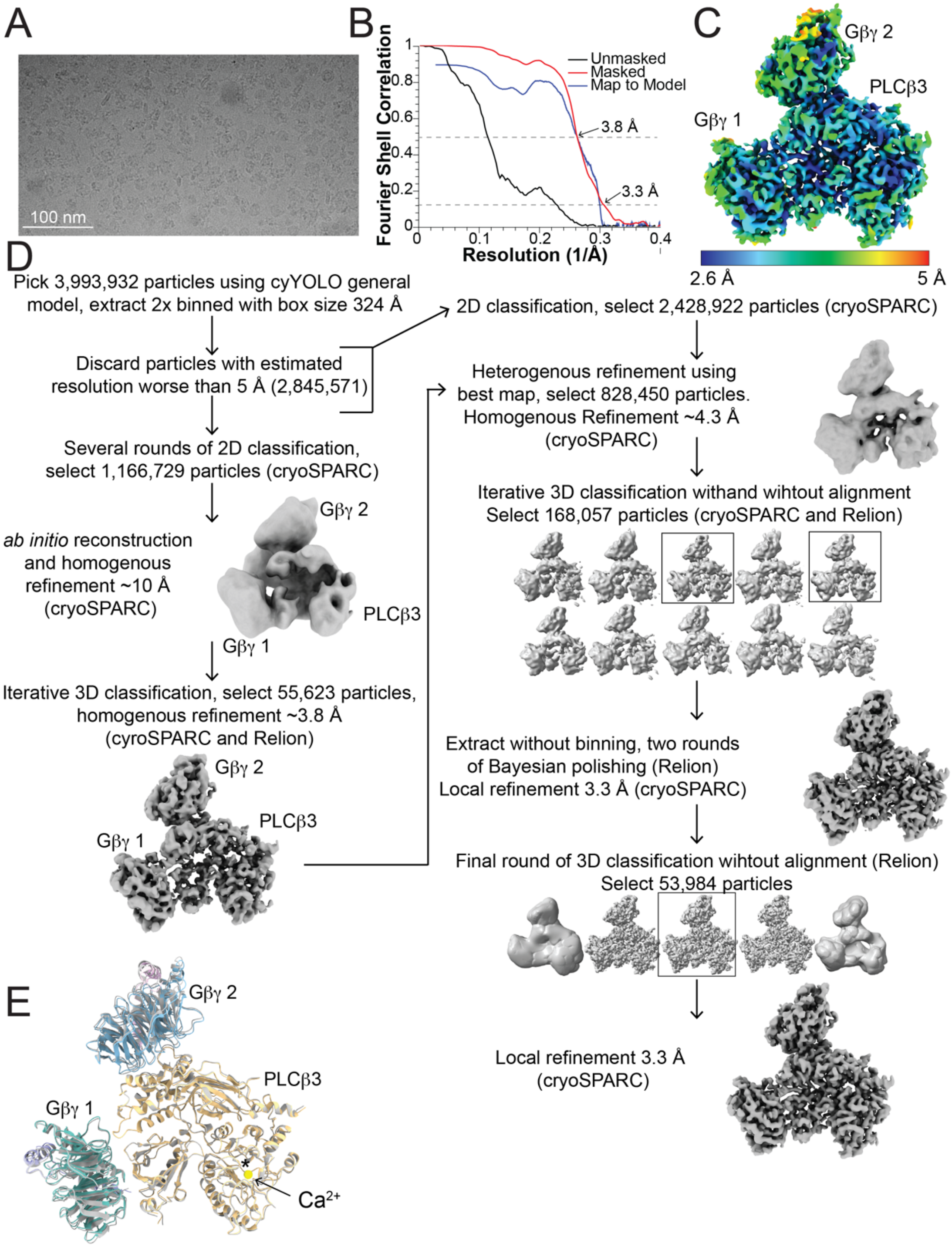


Figure S7 (related to Figure 5, 6): Structure determination of $PLC\beta3 \cdot G\beta\gamma$ complex on nanodiscs A: representative micrograph. B: Fourier shell correlation (FSC) curves for the unmasked (black) and masked (red) maps and between the map and model (blue). The 0.143 and 0.5 thresholds are denoted by dashed lines. C: Final masked, sharpened map colored by local resolution determined by cryoSPARC. D: Summary of data processing steps, see methods. Maps shown are unsharpened. E: Structural alignment of the $PLC\beta3 \cdot G\beta\gamma$ complex on liposomes (colored by protein) and on nanodiscs (gray). $PLC\beta3$ is yellow, $G\beta1$ is dark teal, $G\gamma1$ is light purple, $G\beta2$ is light blue and $G\gamma 2$ is light pink. RMSD ~0.8 Å. Calcium ion from the nanodisc structure is shown as a yellow sphere and the active site is denoted with an asterisks.

### References


1. Ford, C.E., et al., *Molecular basis for interactions of G protein βΓ subunits with effectors.* Science, 1998. **280**(5367): p. 1271-1274.

2. Wagner, T., et al., *SPHIRE-crYOLO is a fast and accurate fully automated particle picker for cryo-EM.* Communications Biology, 2019. **2**(1): p. 1-13.

3. Lyon, A.M., et al., *Full-length Gαq-phospholipase C-β3 structure reveals interfaces of the C-terminal coiled-coil domain.* Nature Structural and Molecular Biology, 2013. **20**: p. 355-362.

4. Jezyk, M.R., et al., *Crystal structure of Rac1 bound to its effector phospholipase C-β2.* Nature Structural & Molecular Biology, 2006. **13**(12): p. 1135-1140.
